# Lifestyle-specific *S*-nitrosylation of protein cysteine thiols regulates *Escherichia coli* biofilm formation and resistance to oxidative stress

**DOI:** 10.1101/2020.09.29.318139

**Authors:** Nicolas Barraud, Sylvie Létoffé, Christophe Beloin, Joelle Vinh, Giovanni Chiappetta, Jean-Marc Ghigo

**Affiliations:** *Genetics of biofilms Laboratory, Institut Pasteur*, UMR CNRS2001, *75015 Paris*, France; SMBP, ESPCI Paris, Université PSL, CNRS, 75005 Paris, France

**Keywords:** biofilms, *S*-nitrosylation, cysteine, *Escherichia coli*, SILAC, redox proteomics

## Abstract

Communities of bacteria called biofilms are characterized by reduced diffusion, steep oxygen and redox gradients and specific properties compared to individualized planktonic bacteria. In this study, we investigated whether signaling via nitrosylation of protein cysteine thiols (*S*-nitrosylation), regulating a wide range of functions in eukaryotes, could also specifically occur in biofilms and contribute to bacterial adaptation to this widespread lifestyle. We used a redox proteomic approach to compare cysteine *S*-nitrosylation in aerobic and anaerobic biofilm and planktonic *Escherichia coli* cultures and we identified proteins with biofilm-specific *S*-nitrosylation status. Using bacterial genetics and various phenotypic screens, we showed that impairing *S*-nitrosylation in proteins involved in redox homeostasis and amino acid synthesis such as OxyR, KatG and GltD altered important biofilm properties, including motility, biofilm maturation or resistance to oxidative stress. Our study therefore revealed that *S*-nitrosylation constitutes a physiological basis underlying functions critical for *E. coli* adaptation to the biofilm environment.

## INTRODUCTION

The formation of surface-attached communities of bacteria embedded in a matrix called biofilms provides a microenvironment preserved from external variations, which allows biofilms to colonize most surfaces, with both positive or negative ecological, medical and industrial consequences (Flemming et al., 2016; Hall-Stoodley et al., 2004). Compared to free-floating, individualized planktonic bacteria, biofilms develop specific metabolic capabilities, including a high tolerance to antimicrobials and host immune defenses (Chalabaev et al., 2014; Jensen et al., 2010; Lebeaux et al., 2014). However, whereas the understanding of bacterial adaptations to the biofilm lifestyle could provide clues for biofilm control, the physiological bases underlying these adaptations are still poorly understood.

One key aspect of the biofilm microenvironment is its physicochemical heterogeneity regarding levels of nutrient, wastes or oxygen (O_2_) (Stewart and Franklin, 2008). Indeed, steep O_2_ gradients were shown to develop rapidly in the 3-dimensional biofilm structure (Barraud et al., 2006; De Beer et al., 1994; Karampatzakis et al., 2017; Wessel et al., 2014) and transcriptomics studies in facultative or strict aerobes such as *Escherichia coli* or *Pseudomonas aeruginosa* revealed gene expression profiles consistent with metabolic adaptations to biofilm microaerobic or anaerobic conditions (Beloin et al., 2004; Harms et al., 2016; Hentzer et al., 2005; Letoffe et al., 2017; Whiteley et al., 2001).

Changes in O_2_ levels in biofilms have important physiological consequences due to alteration of redox conditions and redox-mediated signaling in response to endogenous or exogenous oxidative or nitrosative stress (Bueno et al., 2012; Ezraty et al., 2017; Okegbe et al., 2012; Sporer et al., 2017). Metabolic redox processes often generate reactive oxygen and nitrogen intermediates such as hydroxyl radical (OH·) leading to hydrogen peroxide (H_2_O_2_), or nitric oxide (NO) produced during anaerobic respiration on nitrate (Corker and Poole, 2003; Imlay, 2013). These highly reactive intermediates can, in turn, activate various signaling pathways via covalent binding to protein sensors including cysteine thiols, heme and non-heme metal centers or iron-sulfur clusters (D’Autreaux and Toledano, 2007; Paulsen and Carroll, 2013; Vazquez-Torres, 2012).

Cysteine thiols are of particular interest as they are involved in a range of reversible redox states from reduced S-H (oxidation number −2), disulfide bridges S-S (−1), nitrosylated S-NO (0), or sulfenic acids S-OH (0), associated with profound changes in corresponding protein functions (Conte and Carroll, 2013). However, whereas the formation of disulfide bridges in proteins exported to the periplasmic space or under conditions of oxidative stress is a well-studied process in bacteria (Landeta et al., 2018), much less is known about bacterial *S*-nitrosylation redox signaling. This reversible, enzymatically-controlled modification involves the reaction of a nitrosonium cation NO+ with a reduced thiol (Fernando et al., 2019) and is known to regulate a wide range of cellular functions in plants and animals (Benhar and Stamler, 2005; Hess et al., 2005; Lindermayr et al., 2005), as well as host-microbe interactions, including pathogenesis and defense mechanisms (Bang et al., 2006; Chung et al., 2013; Laver et al., 2010; Potter et al., 2009; Rhee et al., 2005), or synergistic interactions (Seth et al., 2019). However, redox sensors regulated by *S*-nitrosylation have been poorly studied in bacteria, except for the *E. coli* transcriptional regulator OxyR (Green and Paget, 2004). OxyR ability to bind target DNA sequences varies upon conformational changes that depend on its redox state, thus acting as a redox switch for OxyR-dependent transcription (Aslund et al., 1999). Previous studies showed that oxidative stress generated from H_2_O_2_ led to the formation of a disulfide bond between Cys199 and Cys208, activating an oxidative stress response in *E. coli* (Zheng et al., 1998). OxyR is also known to regulate other processes including adhesion and autoaggregation via phase variation regulation of the surface exposed autotransporter adhesin antigen 43, although the role of redox sensing in this signaling remains unclear (Chauhan et al., 2013; Schembri et al., 2003; Wallecha et al., 2003). Finally, recent works suggested that OxyR could adopt additional redox states, including sulfenic S-OH or nitrosylated S-NO, and regulate a specific set of genes under conditions of *S*-nitrosylation (Seth et al., 2012; Seth et al., 2018).

In this study, we hypothesized that specific oxidation and nitrosylation patterns of cysteine thiols may occur in *E. coli* biofilm and not in planktonic conditions, thus contributing to the development of biofilm *S*-nitrosylation signaling and functions. We used a redox proteomics method combining the biotin-switch detection of protein *S*-nitrosylation with Stable Isotope Labeling in Cell Culture (SILAC) to identify proteins with biofilm-specific *S*-nitrosylated cysteine thiols. This approach showed that impairing *S*-nitrosylation status of proteins involved in redox homeostasis and amino acid synthesis affects *E. coli* biofilm formation and oxidative stress resistance, therefore identifying *S*-nitrosylation as a mechanism regulating functions critical for *E. coli* adaptation to the biofilm lifestyle.

## RESULTS

### Development of a biotin-switch protocol to detect protein *S*-nitrosylation in *E. coli* planktonic and biofilm cultures

In order to detect *S*-nitrosylated (S-NO) peptides in *E. coli*, we adapted a previously described biotin-switch approach (Li et al., 2015) based on (i) the addition of iodoacetamide (IAM) to block free, reduced thiols, followed by (ii) the mild and selective reduction of *S*-nitrosylated cysteines with ascorbate and IAM-biotin labeling (Fig. 1A). The extent of protein *S*-nitrosylation was then assessed by western blotting and immunodetection, taking advantage of avidin-biotin affinity with avidin-HRP antibodies.

**Figure 1.**
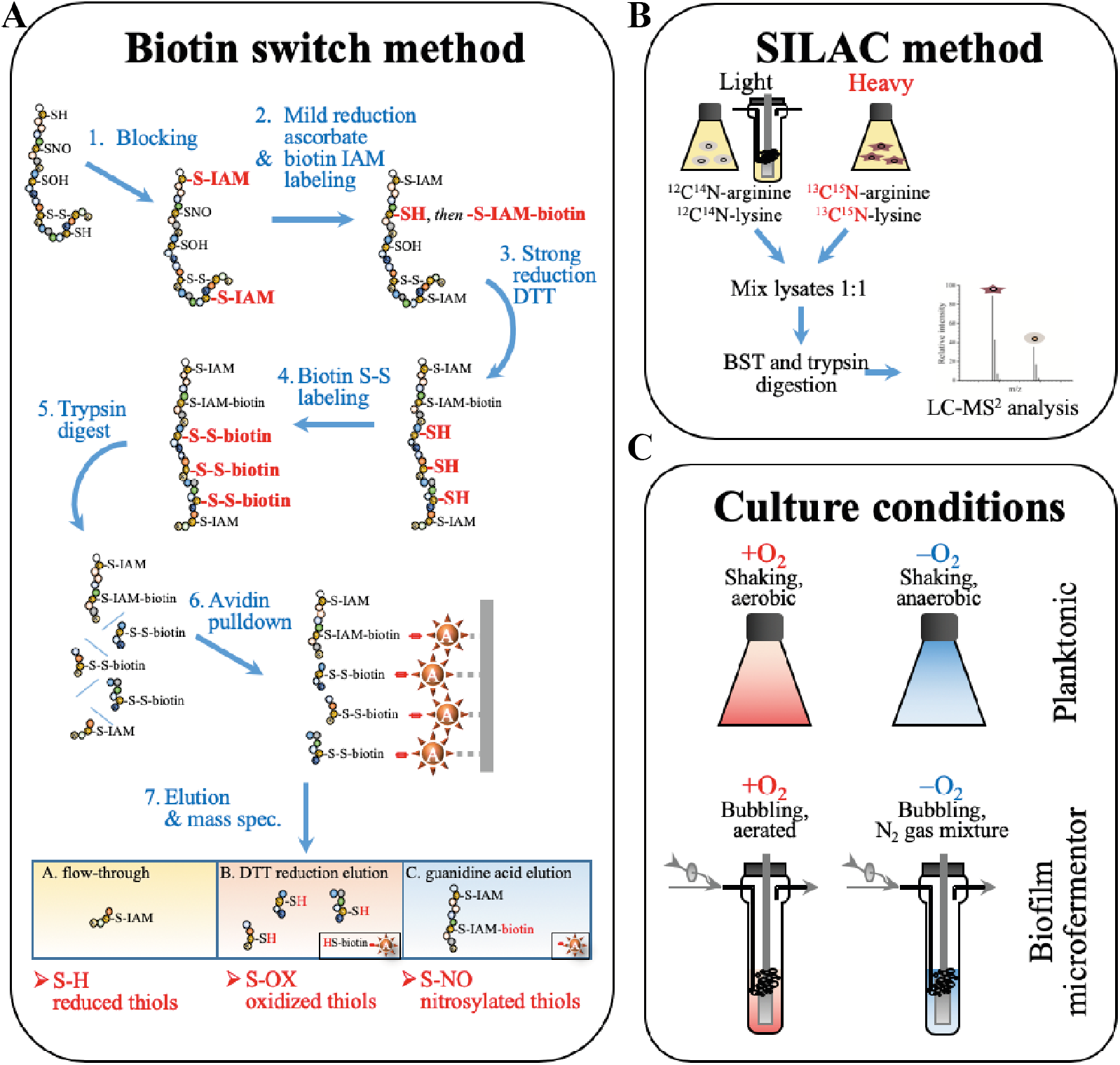
Workflow of the biotin-switch SILAC method used to identify and quantify redox-modified protein thiols in *E. coli* biofilm and planktonic conditions. **A:** The biotin-switch method consists of the sequential blocking, reduction, and labeling of cysteine thiols; for the differential detection of S-NO and S-OX, cysteines are first reduced with ascorbate, a mild reducing agent, and then with the strong reducing agent DTT. Labeled peptides are then selectively eluted and collected for identification by mass spectrometry (LC-MS/MS). **B:** Accurate peptide quantification using a SILAC approach was performed on each sample using heavy isotope lysine and arginine labeled samples as internal references. **C:** *E. coli* proteins were extracted from 4 different cultures: planktonic shake flasks or biofilm microfermentors, each under aerobic or anaerobic conditions. Each condition included 5 biological replicates.

We validated the initial steps of the biotin-switch protocol using planktonic *E. coli* cultures grown in anaerobic conditions, either in the presence of excess amounts of nitrate, which leads to the accumulation of *S*-nitrosylated proteins (S-NO+ conditions), or in the presence of fumarate, preventing *S*-nitrosylation (S-NO-conditions) (Seth et al., 2012). Total proteins extracted from 24 h cultures were processed following the biotin-switch protocol, except that some samples were either (i) not blocked with IAM, or (ii) not reduced with ascorbate. In the absence of IAM, cysteines were biotinylated in all conditions (Fig. 2 lanes a,b). In contrast, the use of IAM blocking agent without subsequent reduction with ascorbate led to minimal biotin-labeled cysteine signal in both S-NO+ and S-NO-conditions (Fig. 2 lanes c,d), potentially corresponding to naturally biotinylated proteins. Finally, when protein cysteine thiols were blocked with IAM and S-NO specifically reduced with ascorbate, S-NO+ samples (nitrate conditions) showed abundant cysteine labeling (Fig. 2 lane e), while S-NO-samples (fumarate conditions) did not (Fig. 2 lane f). This indicated that the use of IAM and ascorbate could selectively label *E. coli S*-nitrosylated proteins in our experimental conditions.

**Figure 2.**
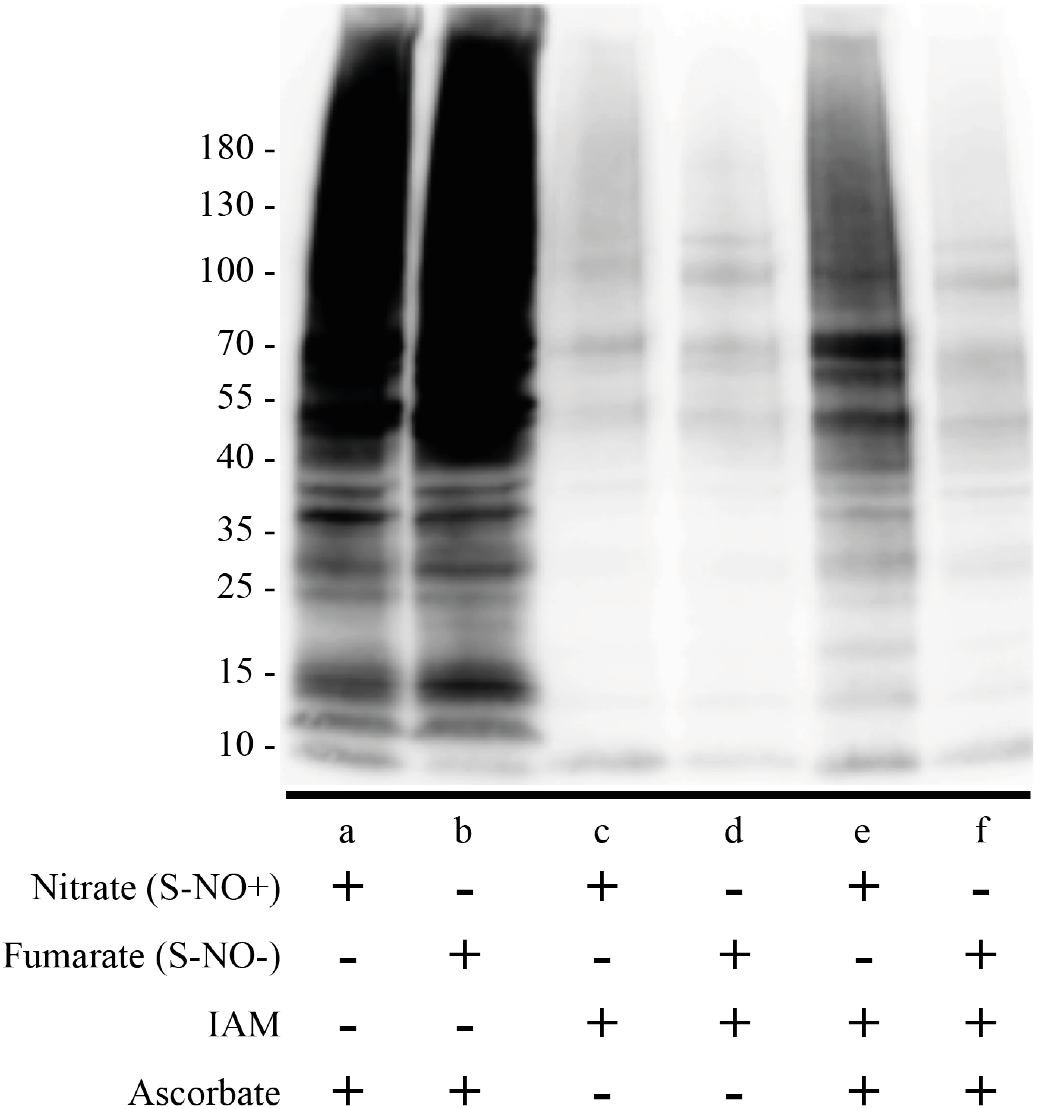
Validation of the biotin-switch method to detect protein *S*-nitrosylation in *E. coli*. Whole proteins were extracted from *E. coli* planktonic cultures grown under *S*-nitrosylating (S-NO+, anaerobic with excess nitrate), or non-*S*-nitrosylating conditions (S-NO-, anaerobic with fumarate). Proteins were then processed following the biotin-switch technique, except that some samples were not blocked with IAM (lanes a,b), or not reduced with ascorbate (lanes c,d), before labeling with IAM-biotin and immunodetection using avidin-HRP antibodies.

To study potential modifications of protein *S*-nitrosylation profile in biofilms, we used continuous-flow biofilm microfermentors to grow mature *E. coli* biofilms in aerobic and anaerobic conditions in M63B1 glucose minimal medium supplemented with nitrate (S-NO+ conditions) or fumarate (S-NO-conditions) and we compared to the S-NO profiles of corresponding planktonic aerobic or anaerobic cultures (Fig. 1C). Biofilms grown in absence of O_2_ and under conditions favoring *S*-nitrosylation (+ nitrate) did not display more S-NO biotin-labeled signals compared to corresponding planktonic bacteria. However, biofilm S-NO labeled samples all displayed distinct band patterns, suggesting specific protein *S*-nitrosylation in biofilm bacteria (Fig. 3 compare lanes a and c). Furthermore, planktonic cultures showed increased *S*-nitrosylation when grown under aerobic compared to anaerobic conditions (Fig. 3 compare lanes c and g), whereas growing biofilm grown under high aeration did not drastically alter protein *S*-nitrosylation profile (Fig. 3, compare lanes a and e). Taken together, these results indicated that reduced availability of O_2_ availability in biofilms or anaerobic conditions led to reduced levels of *S*-nitrosylation.

**Figure 3.**
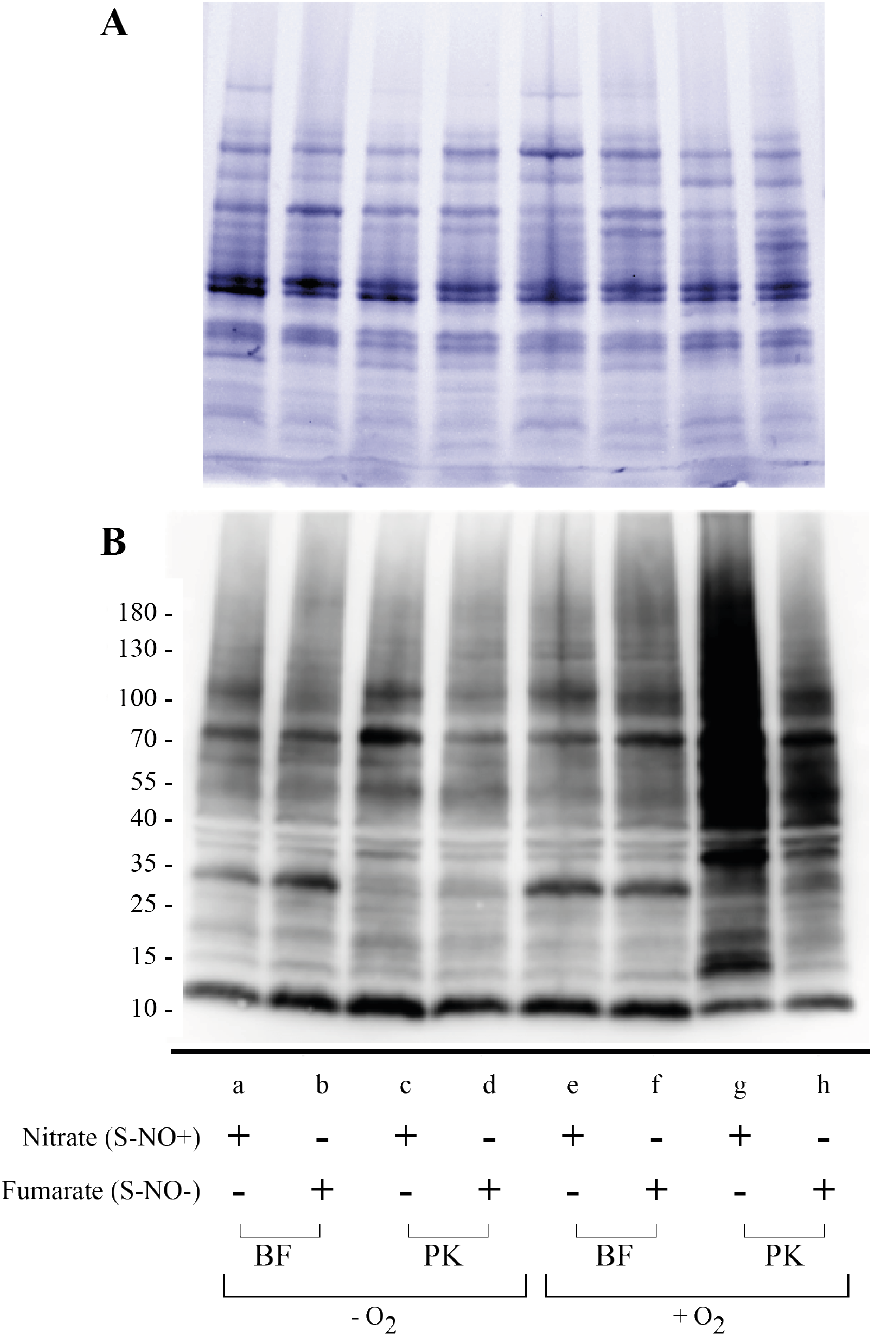
Protein *S*-nitrosylation profiles in *E. coli* planktonic (PK) or biofilm (BF) conditions. Cultures were grown under aerobiosis (+O_2_) or anaerobiosis (−O_2_) and under *S*-nitrosylating conditions with excess 10 mM nitrate (S-NO+), or non-*S*-nitrosylating condition supplemented with 10 mM fumarate (S-NO-). Proteins were extracted, processed with the biotin-switch method (see Fig. 1) and analyzed by western blot. **A:** Total protein extracts visualized with Stain-free fluorescent detection; **B:** S-NO proteins visualized by western blot followed by immunodetection using avidin-HRP antibodies.

### Redox proteomics analysis reveals fewer S-OX and S-NO cysteines but more reduced cysteines in biofilms compared to planktonic cultures

In order to identify proteins with cysteines specifically *S*-nitrosylated in biofilm and planktonic conditions, we combined our biotin-switch method with previously described redox proteomics workflow using Stable Isotope Labeling in Cell Culture (SILAC) (Fig. 1B) (Li et al., 2015). Each sample contained equal amounts of proteins originating from isogenic arginine and lysine *E. coli* auxotroph strains grown in presence of the stable isotope-labeled (^13^C^15^N) L-arginine and L-lysine forms. In addition to the labeling of S-NO cysteines with IAM-biotin, reversibly oxidized cysteines (S-OX), including disulfide bridges and sulfenic acids, were labeled with biotin-HPDP. Hence, three peptide fractions were obtained after affinity enrichment: (i) the unbound (reduced cysteines and cysteine-free peptides), (ii) the DTT fraction (S-OX) and (iii) the guanidine fraction (S-NO) (Fig. 1A bottom). Overall, we quantified the level of 1426 proteins (854 and 892 in aerobic and anaerobic conditions, respectively), 179 S-NO cysteines (161 aerobic; 132 anaerobic), 410 S-OX cysteines (357 aerobic; 249 anaerobic), and 386 reduced cysteines (243 aerobic-Supplementary Table S1; 341 anaerobic - Supplementary Table S2). Volcano plot visualization of the data showed that the distributions of proteins fold changes of proteins and reduced cysteine were symmetric and overlapped between the two studied bacterial lifestyles (Supplementary Figure S1). By contrast, the distributions of S-OX and S-NO cysteines deviated towards the planktonic mode of life, suggesting potential changes in cysteine site occupancy between the two bacterial lifestyles (Olsen et al., 2010). According to site occupancy equations (Olsen et al., 2010), these trends suggested that the detected S-OX and S-NO cysteines involved a small fraction of the associated proteins.

Consistent with our preliminary immunodetection analysis (Fig. 3), we identified more cysteines significantly *S*-nitrosylated in planktonic than in biofilm bacteria (38 *vs*. 6, see Tables 1 and 2). Under aerobic conditions, 26 cysteines (corresponding to 25 proteins) showed increased nitrosylation levels in planktonic bacteria compared to biofilms, while only two S-NO cysteines were more abundant in biofilms compared to planktonic bacteria (Table 1). Similarly, under anaerobic conditions, 12 cysteines (11 proteins) had increased nitrosylation levels in planktonic cells, while 4 S-NO cysteines were more abundant (Table 2). Furthermore, we identified more S-NO cysteines in cells grown under aerobic conditions, with 28 cysteines in biofilm and planktonic cells, compared to 16 cysteines in anaerobic biofilm and planktonic cells.

**Table 1.**
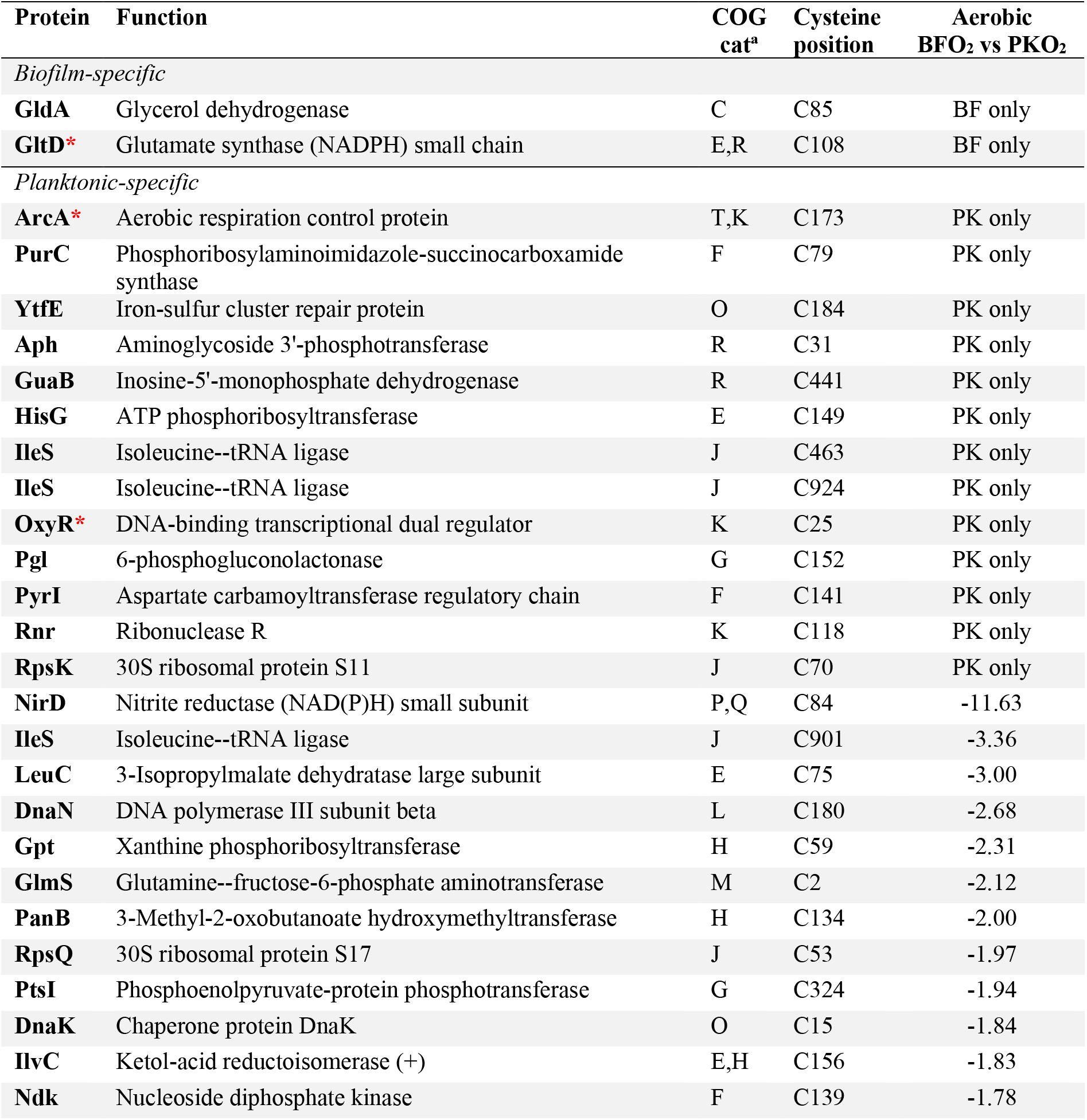
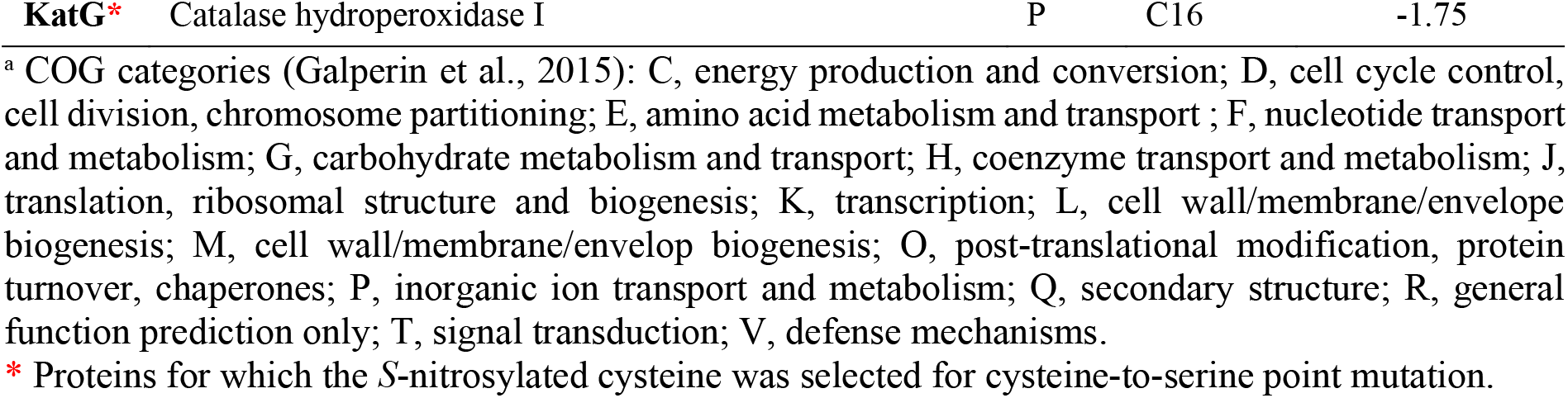
List of cysteine residues identified to be differentially *S*-nitrosylated in aerobically-grown biofilm *vs* planktonic *E. coli* cultures by using the biotin-switch SILAC method. PKO_2_, planktonic with O_2_; BFO_2_, biofilm with O_2_. Post-translationally modified peptides that were detected only in one type of sample are listed as “BF only” (biofilm) or “PK only” (planktonic). Peptides with significant changes in S-NO profile with a p-value <0.05 are shown (n=5); data indicate fold-changes in modified peptide normalized to total peptide count for biofilm compared to planktonic samples.

**Table 2.** List of protein cysteines identified to be differentially *S*-nitrosylated in anaerobically grown biofilm *vs* planktonic *E. coli* cultures by using the biotin-switch SILAC method. PKn, planktonic without O_2_; BFn, biofilm without O_2_. Post-translationally modified peptides that were detected only in one type of sample are listed as “BF only” (biofilm) or “PK only” (planktonic). Peptides which showed significant changes in S-NO profile with a p-value <0.05 are shown (n=5); data indicate fold-changes in modified peptide normalized to total peptide count for biofilm compared to planktonic samples.

| Protein | Function | COG cat<sup>a</sup> | Cysteine position | Anaerobic BF<sub>n</sub> vs PK<sub>n</sub> |
| --- | --- | --- | --- | --- |
| *Biofilm-specific* |  |  |  |  |
| **GrxC\*** | Glutaredoxin 3 | O | C66 | BF only |
| **LeuD\*** | 3-Isopropylmalate dehydratase small subunit | E | C82 | BF only |
| **LuxS\*** | S-ribosylhomocysteine lyase | T | C41 | 94.56 |
| **IlvE\*** | Branched-chain-amino-acid aminotransferase | EH | C42 | 38.42 |
| *Planktonic-specific* |  |  |  |  |
| **AroG** | Phospho-2-dehydro-3-deoxyheptonate aldolase | E | C208 | PK only |
| **LuxS\*** | S-Ribosylhomocysteine lyase | T | C128 | PK only |
| **RplJ** | 50S ribosomal protein L10 | J | C71 | PK only |
| **OxyR\*** | DNA-binding transcriptional dual regulator | K | C25 | PK only<sup>#</sup> |
| **AtpD** | ATP synthase subunit beta | C | C138 | -6.66 |
| **Tpx** | Thiol peroxidase | O | C95 | -3.41 |
| **PykA** | Pyruvate kinase II | G | C200 | -2.58 |
| **AhpC** | Alkyl hydroperoxide reductase C | V | C166 | -2.30 |
| **KatG\*** | Catalase hydroperoxidase I | P | C16 | -1.95 |
| **GroL** | Chaperonin | O | C138 | -1.88 |
| **GroL** | Chaperonin | O | C458 | -1.79 |
| **Lrp** | Leucine-responsive regulatory protein | K | C45 | -1.76 |

<sup>a</sup> COG categories (Galperin et al., 2015) as in Table 1.
\* Proteins for which the *S*-nitrosylated cysteine was selected for cysteine-to-serine point mutation.
<sup>#</sup> OxyR-C25-S-NO was detected in only 3 replicates in anaerobically grown planktonic cells and in none of the anaerobically grown biofilm samples (as indicated in Materials and Methods, the selection threshold used in this study was 4 detections in one condition and none in the other).

Our redox proteomics protocol also allowed us to analyze S-OX cysteines (engaged in disulfide bonds or other reversible oxidation states), showing, similarly to S-NO cysteines, higher S-OX cysteine levels in planktonic conditions (Supplementary Figure S1). Under aerobiosis, decreased S-OX levels of 61 cysteines (52 proteins) were detected in biofilms compared to planktonic bacteria, while only four cysteines showed increased S-OX levels. Moreover, 11 peptides showed decreased levels of reduced cysteines and five peptides showed increased levels of reduced cysteines (Supplementary Table S1). A similar trend was observed in cultures grown under anaerobic conditions, with 66 detected cysteine residues (57 proteins) with decreased S-OX levels and nine residues (8 proteins) with higher S-OX levels in biofilms compared to planktonic cells. By contrast biofilm showed 14 and 16 cysteine residues displaying increased or decreased levels of cysteine reduction, respectively (Supplementary Table S2). Overall, these data indicated a higher level of cysteine oxidation in planktonic and aerobic conditions compared to biofilm cells.

### Lifestyle-specific *S*-nitrosylated proteins are involved in redox homeostasis and amino acid synthesis

Several proteins involved in redox homeostasis control were differentially *S*-nitrosylated in biofilms compared to planktonic bacteria. Under anaerobic conditions, GrxC was more *S*-nitrosylated in biofilms, while Tpx and AhpC were more *S*-nitrosylated in planktonic cells (Table 2). Under aerobic conditions, the nitrite reductase NirD involved in anaerobic respiration and ArcA, a regulator of aerobic respiration, were also more *S*-nitrosylated in planktonic cells (Table 1). Interestingly, the catalase hydroperoxidase II KatG and the major dual redox regulator OxyR appeared to be *S*-nitrosylated in planktonic bacteria under both aerobic and anaerobic conditions. These results were consistent with varying oxidative conditions in biofilms and the associated presence of gradients of O_2_ or other electron acceptors. Surprisingly, the change in S-NO modification for OxyR was on Cys25, a thiol that had not been identified as *S*-nitrosylated in previous studies using planktonic *E. coli* (Kim et al., 2002; Seth et al., 2012; Seth et al., 2018). Interestingly 2 AhpC cysteines, Cys47 and Cys166 were more oxidized in anaerobic planktonic conditions (Supplementary Table S2), suggesting a potential swap between S-NO and S-OX states, depending on the bacterial lifestyle.

Several of the most highly *S*-nitrosylated peptides in biofilms were involved in amino acid synthesis including GltD under aerobic conditions and LeuD and IlvE under anaerobic conditions. By contrast, LeuC, IlvC and IleS were specifically *S*-nitrosylated in aerobic planktonic bacteria. Interestingly, Lrp, which is involved in the regulation of branched chain amino acids Leu, Ile and Val, was found to be nitrosylated in anaerobic planktonic bacteria on Cys45 (Table 2). This cysteine corresponds to the start of the DNA binding helix-turn-helix domain (Hart et al., 2011), suggesting a potential impact on its regulatory effect. Finally, LuxS, a central element of the quorum sensing cell-cell signaling system, showed a transfer of nitrosylation site from Cys128 in planktonic to Cys41 in biofilms under anaerobic conditions (Fig. 4).

**Figure 4.**
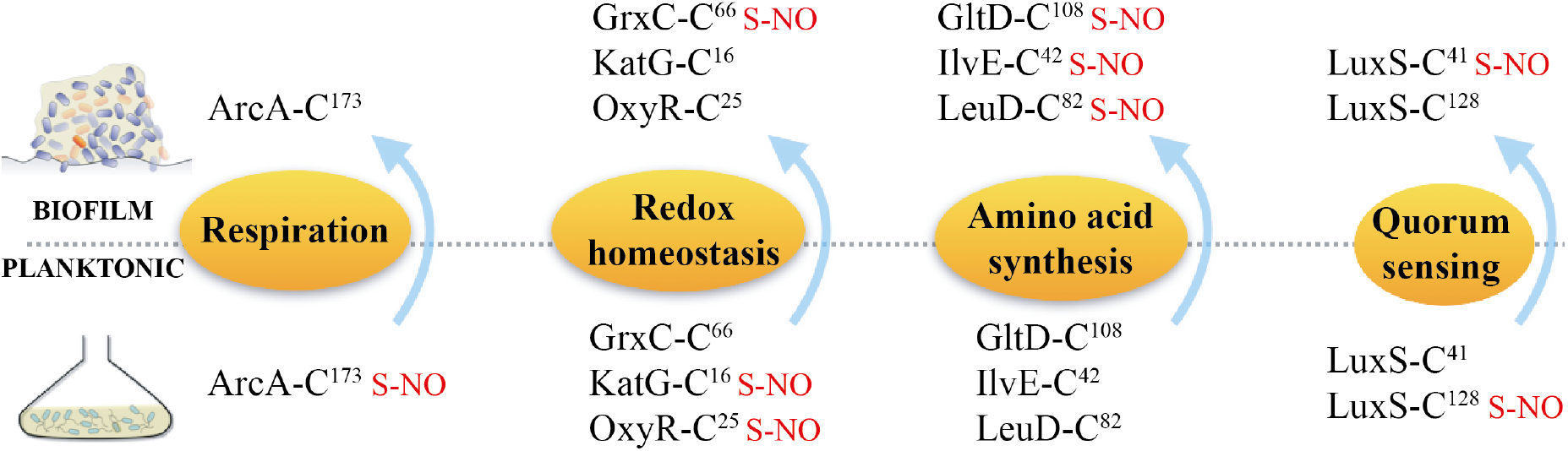
Cysteines with the highest *S*-nitrosylation fold-changes or differentially *S*-nitrosylated in both aerobic and anaerobic conditions. These cysteines are in proteins involved in known biofilm associated phenotypes, including respiration, redox homeostasis, amino acid synthesis and quorum sensing.

### Impairing *S*-nitrosylation status of OxyR, KatG and GltD affects biofilm-associated phenotypes

To investigate the phenotypic consequences of *S*-nitrosylation, we selected proteins displaying high cysteine *S*-nitrosylation fold-changes in biofilm (GltD-Cys108, GrxC-Cys66, LeuD-Cys82, IlvE-Cys42, LuxS-Cys41) or planktonic conditions (ArcA-Cys173, LuxS-Cys128), or that were found to be S-nitrosylated in both anaerobic and aerobic planktonic conditions (OxyR-Cys25 and KatG-Cys16). These cysteines did not have life-style associated modification of their oxidation or reduction status, suggesting a specific *S*-nitrosylation biofilm pattern (Supplementary Tables S1 and S2). We then introduced chromosomal and markerless point mutations changing the codon corresponding to the identified S-NO cysteine by a serine codon (C to S mutation). Cysteine to serine mutations in GrxC (C66S), LeuD (C82S), IlvE (C42S), LuxS (C41S and C128S) or ArcA (C173S) had no detectable impact on growth rate, biofilm formation, H_2_O_2_ sensitivity or motility (Supplementary Fig. S2). We also showed that a mutant in the hybrid cluster protein Hcp previously found to regulate protein *S*-nitrosylation and mediate bacterial motility under conditions of *S*-nitrosylation has no impact on biofilm formation (Sup Fig S3) (Seth et al., 2018). By contrast, *oxyR*_C25S_, *katG*_C16S_ and *gltD*_C108S_ mutants displayed growth rates similar to the WT (Supplementary Fig. S2), but showed increased biofilm formation compared to WT, a phenotype observed both in rich (Fig. 5A) and minimal medium (Supplementary Fig. S4), and which could be complemented by introducing the corresponding plasmid-based WT allele of *oxyR*, *katG* or *gltD* into the corresponding mutant strains (Fig. 5A).

**Figure 5.**
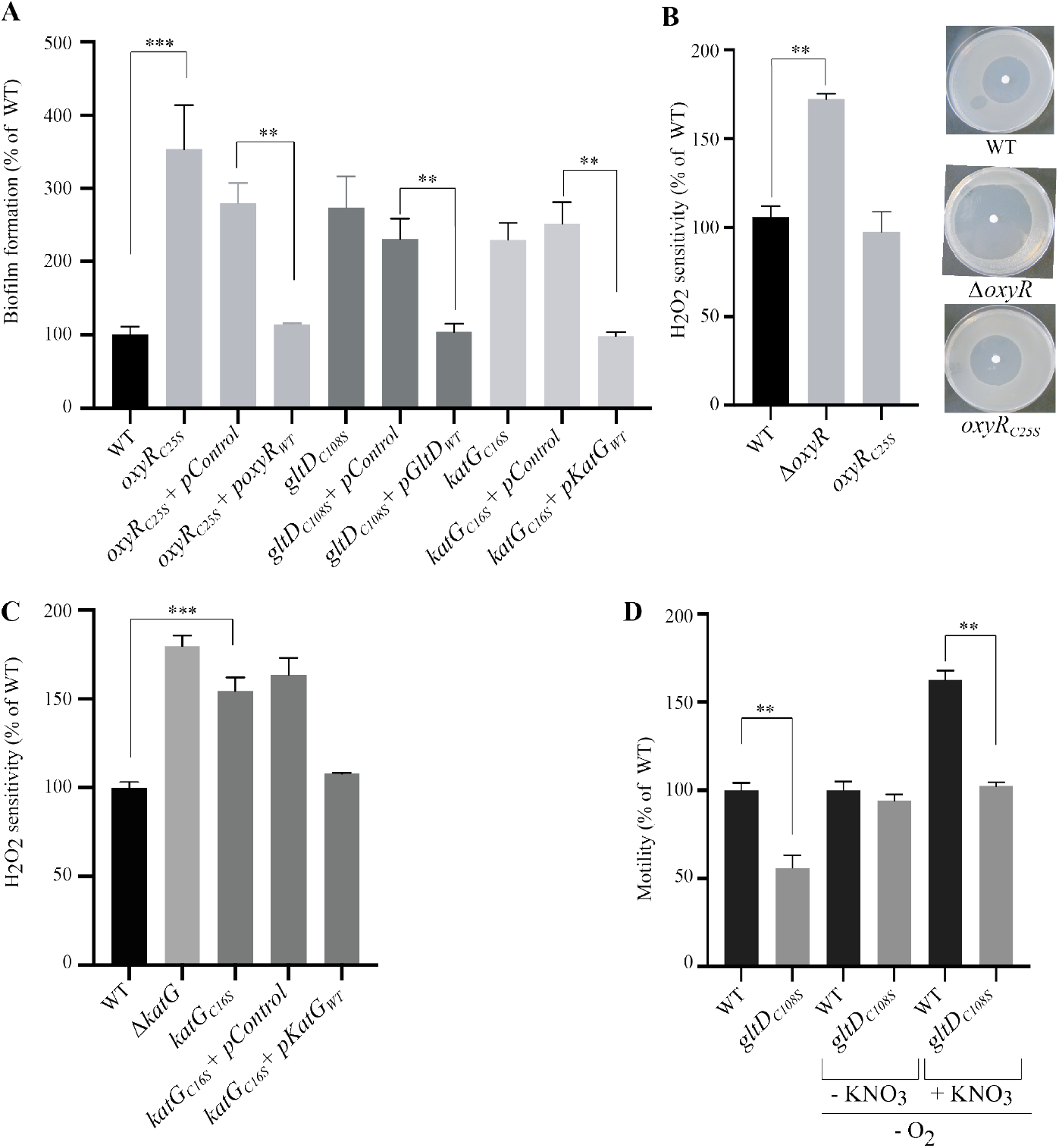
Impairing *S*-nitrosylation status of OxyR, KatG and GltD affects biofilm-associated phenotypes. **A**: Biofilms of *E. coli* WT, *oxyR*_C25S_, *katG*_C16S_ and *gltD*_C108S_ mutants, complemented or not with the corresponding plasmid-based allele or empty vector, were grown in continuous-flow microfermentors for 24 h in LB medium before quantifying the biofilm biomass. The level of biofilm formed by the WT strain was set to 100%. **B**: Sensitivity to H_2_O_2_ oxidative stress of an *oxyR*_C25S_ mutant compared to the WT and Δ*oxyR* mutant. The distance of growth inhibition from the edge of the disk to the edge of the growth zone was measured and was set to 100% for the WT strain. **C**: Increased sensitivity to oxidative stress of *katG*_C16S_ mutant. The sensitivities to H_2_O_2_ of *E. coli* WT and *katG*_C16S_ mutant complemented with corresponding plasmid-based allele or empty vector were compared. The distance from the edge of the disk to the edge of the growth zone was measured and was set to 100% for the WT strain. **D**: *S*-nitrosylation-dependent decreased motility of *gltD*_C108S_ mutant. The motility of *E. coli* WT and *gltD*_C108S_ mutant were compared in aerobic conditions and in anaerobic conditions in presence of KNO_3_. Assays were performed on 0.3% agar plates and incubated overnight at 30°C. All experiments were performed in triplicate, mean values are reported and error bars represent standard deviations. ** p ≤ 0.05, *** p ≤ 0.01. See also Figures S2, S4 and S5.

The cysteines differentially *S*-nitrosylated in biofilms in OxyR, KatG or GltD did not show any biofilm-specific reversible oxidation or reduction (Supplementary Table S1 and Supplementary Table S2), except for OxyR-Cys25, which was more reduced in anaerobic planktonic conditions. This suggested that the altered phenotypes associated with the C to S mutations were due to altered *S*-nitrosylation rather than due to another modification of the redox status of these thiols. The strain carrying the *oxyR*_C25S_ mutation also showed increased cell aggregation, similar to the one of a Δ*oxyR* strain, in which phase-variable expression of Ag43 autotransporter self-aggregation adhesin is locked ON (Supplementary Fig. S5A) (van der Woude and Henderson, 2008). Consistently, *E. coli oxyR*_C25S_ biofilm phenotype depended on the presence of *flu*, the gene encoding Ag43 adhesin (Supplementary Fig. S5B), displayed 100% ON colonies (Supplementary Fig. S5C), and an increased level of Ag43 protein, similarly to a Δ*oxyR* mutant (Supplementary Fig. S5D). Although this suggested that preventing Cys25 *S*-nitrosylation phenocopied an *oxyR* deletion, we did not observe the characteristic increased sensitivity to H_2_O_2_ oxidative stress, indicating that preventing OxyR *S*-nitrosylation affected some but not all OxyR functions (Fig. 5B). To assess the specificity of C25S mutation, we also tested a Q29S mutant potentially impairing OxyR DNA binding site and found that this strain had similar phenotypes as the C25S mutant: increased biofilm formation and no impact on H_2_O_2_ sensitivity (Sup Fig S6). In contrast to *oxyR*_*C*25S_, the *katG*_C16S_ mutant exhibited increased sensitivity to oxidative stress upon exposure to H_2_O_2_ compared to WT (Fig. 5C and Supplementary Fig. S4), showing that planktonic *S*-nitrosylation could protect against oxidative stress. On the other hand, whereas the *gltD*_C108S_ mutation had no impact on sensitivity to oxidative stress (Supplementary Fig. S2), its increased biofilm formation capacity corresponded with a reduced motility phenotype in aerobic condition and in anaerobic condition with KNO_3_ (S-NO+ conditions), indicating that GltD-Cys108 *S*-nitrosylation increases motility in biofilm bacteria (Fig. 5D and Supplementary Fig. S4). Taken together, these results showed that altering *S*-nitrosylation can trigger an array of functions relevant to the switch between the biofilm and planktonic lifestyles.

## DISCUSSION

Bacterial biofilms are characterized by steep O_2_ gradients leading to microaerobic or anaerobic zones, profoundly affecting the physiology of biofilm bacteria. In this study, we used redox proteomics and bacterial genetic and phenotypic analyses to identify nitrosylated cysteines altering biofilm functions.

The addition of a reduction and labeling step to a previously described *S*-oxidized (S-OX) cysteine biotin-switch protocol (Li et al., 2015), combined with SILAC allowed us to selectively detect and quantify *S*-nitrosylated (S-NO) cysteines and other redox protein modifications in *E. coli* biofilm and planktonic bacteria. We showed that proteins extracted from planktonic bacteria possess more S-NO and S-OX cysteines compared to protein extracted from poorly oxygenated biofilm environment. Furthermore, increased *S*-nitrosylation of proteins in the presence of O_2_ was proposed to increase likelihood of NO moieties binding to cysteine thiols (Foster and Stamler, 2004), which is consistent with higher S-NO and S-OX levels detected in aerobiosis. Our approach allowed us to monitor global changes in protein expression profiles, comparing biofilms and planktonic cultures. Consistently with previous standard proteomic analyses (Collet et al., 2008), the most downregulated proteins in aerobic biofilms corresponded to siderophore biosynthesis and iron transport (Supplementary Figure S7). In contrast, upregulated proteins in aerobic biofilms were clustered into respiration, carbohydrate uptake, or amino acids biosynthesis functional class (Supplementary Figure S7).

Our study showed that reversible redox modifications of proteins previously associated with the biofilm lifestyle, including redox homeostasis, amino acid synthesis, or respiration, occur during biofilm development. While these protein modifications may be a consequence of the biofilm reducing environment, the presence of biofilm-specific S-NO proteins also suggest that *S*-nitrosylation-dependent regulation could control biofilm functions. Recently, a multiplex enzymatic mechanism was identified for the regulation of cell motility and metabolism in *E. coli*, involving the hybrid cluster protein Hcp and nitrate reductase NarG. Under anaerobic conditions, Hcp was found to interact with several proteins to induce S-NO from NarG-derived NO as well as propagate S-NO-based signaling via trans-nitrosylation of proteins (Seth et al., 2018). However, when we tested a Δ*hcp* mutant, no impact on biofilm formation was observed, suggesting that Hcp is not involved in the regulation of S-NO dependent biofilm formation.

Protein *S*-nitrosylation is known to regulate a wide range of critical physiological functions in eukaryotes (Hess et al., 2005; Smith and Marletta, 2012). Here, we demonstrated that disabling *S*-nitrosylation sites in OxyR, KatG or GltD increases biofilm development. Since KatG-Cys16 cysteine is *S*-nitrosylated in planktonic conditions under both aerobic and anaerobic conditions, this suggests that KatG *S*-nitrosylation is associated with the switch from biofilm to planktonic lifestyle, either by directly inhibiting bacterial adhesion mechanisms and promoting biofilm dispersal or via an intermediate regulator. Furthermore, the increased sensitivity of *katG*_C16S_ mutant towards oxidative stress also indicates a role for KatG-Cys16 *S*-nitrosylation in the regulation of oxidative and nitrosative stress in *E. coli* (Adolfsen et al., 2019). In contrast to KatG, GltD-Cys108 was found to be *S*-nitrosylated in biofilm conditions. Taken together these results indicate that both denitrosylation (*e.g*. KatG) and nitrosylation (*e.g*. GltD) mechanisms may be involved in biofilm formation.

OxyR is a well-known regulator of the phase variable switch of the autotransporter adhesin Ag43, a major *E. coli* surface protein involved in bacterial aggregation and biofilm formation. OxyR contains 6 cysteines, and Cys25 and Cys199 are the only solvent-accessible sites in the native protein (Kim et al., 2002). Previous investigation of NO binding to OxyR cysteines by exposing His-tag purified OxyR to 20-fold molar excess of the NO donor GSNO (Seth et al., 2012) showed that Cys199 was predominantly *S*-nitrosylated in presence of 10 mM nitrate. However, the authors reported that Cys25 also showed low 20% S-NO signal, which they attributed to incomplete thiol blocking during a preliminary step (Seth et al., 2012). Here we found that OxyR is *S*-nitrosylated on Cys25 in planktonic bacteria in the presence of 300 μM KNO_3_, which represents physiologically relevant levels of nitrate (Palmer et al., 2007; Winter et al., 2013), suggesting that Cys25 is a *bona fide* OxyR *S*-nitrosylation site. Furthermore, inactivating this S-NO site led to increased biofilm formation, similar to a Δ*oxyR* mutant. By contrast, OxyR-Cys25 denitrosylation had no impact on *E. coli* sensitivity to oxidative stress, which is typically increased in absence of OxyR (Aslund et al., 1999; Seth et al., 2020). Cys25 is located in the OxyR DNA binding site and the increased expression of Ag43 and bacterial aggregation and biofilm formation could be a consequence of the altered OxyR DNA-binding site in the *oxyR_C25S_* mutant. However, previous studies showed that cysteine-to-serine mutation of Cys25 did not affect binding to the *oxyS* promoter region (Kullik et al., 1995). Moreover, an *oxyR_Q29S_* mutant, potentially impairing OxyR DNA binding site, also increased biofilm formation and had no impact on H_2_O_2_ sensitivity, which is consistent with the observed lack of effect of this mutation on resistance to H_2_O_2_. While these observations do not exclude that NO binding to Cys25 could affect Ag43 gene transcription, we cannot distinguish between phenotypes associated to removal of *S*-nitrosylation and impairment of DNA binding.

Overall this study revealed that proteins involved in functions broadly associated to biofilm physiology across various bacterial species are specifically *S*-nitrosylated in biofilms. The role played by *S*-nitrosylation in the regulation of biofilm development suggests that effectors of S-NO proteins could constitute new targets for biofilm control strategies. How these mechanisms are coordinated in time and space during biofilm development and whether specific enzymes such as nitrosylases or denitrosylases are involved remain to be elucidated.

## STAR METHODS

### Bacterial strains and culture media

Bacterial strains used in this study are listed in supplementary Table S4. Bacterial planktonic cultures were grown in Lysogeny broth (LB) containing 1% (w/v) tryptone, 0.5% (w/v) yeast extract and 1% (w/v) NaCl at 37°C, supplemented with appropriate antibiotics when needed (kanamycin 50 μg/mL, carbenicillin 100 mg/mL, tetracycline 7.5 μg/mL). For biofilm and planktonic comparisons, cultures were grown in minimal medium M63B1 containing 100 mM KH_2_PO_4_, 15 mM (NH_4_)_2_SO_4_, 0.4 mM MgSO_4_, 10 μM FeSO_4_, 3 μM vitamin B1, pH 7.0, and supplemented with 0.1% (w/v) (biofilm) or 0.4% (planktonic) glucose (22 mM). To obtain proteins that were analyzed by mass spectrometry (SILAC), all biofilm and planktonic cultures were supplemented with 300 μM KNO_3_, which represents physiologically relevant levels of nitrate (Palmer et al., 2007; Winter et al., 2013). In contrast, for western-blot analysis (biotin-switch method confirmation), biofilm and planktonic cultures were grown in the presence of 10 mM KNO_3_ or fumarate (excess amounts), as described previously to generate *S*-nitrosylation profiles detectable with this analysis (Seth et al., 2012). For redox proteomics experiments, planktonic and biofilm cultures of auxotroph mutants were supplemented with 500 μM of L-arginine and L-lysine; or for the reference samples, planktonic cultures were supplemented with 500 μM of stable isotopes ^13^C_6_^15^N_4_ L-arginine and ^13^C_6_^15^N_2_ L-lysine (Pierce). All media and chemicals were purchased from Sigma-Aldrich or from specific suppliers as indicated.

### Single nucleotide mutant construction

Single nucleotide mutations were introduced in MG1655 F’*tet* genome using transient mutator multiplex automated genome engineering (TM-MAGE) (Gallagher et al., 2014). Primers consisting of 90 bp long ssDNA fragments harboring the targeted nucleotide mismatch and phosphorothioate bonds at the 5’ end were designed using the online MODEST tool available at http://modest.biosustain.dtu.dk (Bonde et al., 2014) and ordered from Sigma (see Table S5). First, λ-Red- and *dam*-containing plasmid pMA7SacB (Lennen et al., 2016) was transformed into *E. coli*. pMA7SacB-containing *E. coli* induced with 0.2% (w/v) L-arabinose and made electrocompetent by rinsing several times in cold deionized water were repeatedly transformed with 10 pmol of the specific primer (Wang and Church, 2011), and immediately regrown in LB with tetracycline and carbenicillin antibiotics. After 3-4 cycles, single clones were isolated and screened by PCR using a high discrimination HiDi polymerase (myPols) and primers with the targeted original or mutated nucleotide at the 3’ end (see Table S5). Finally, positive clones were checked by PCR with specific primers and DNA sequencing.

### Transduction of chromosomal mutation

Single nucleotide mutations were moved into naïve WT (MG1655 F’*tet*) background by introducing for each of these three strains, a kanamycin resistance marker immediately downstream of the mutated *oxyR, katG* and *gltD* genes. We then performed a P1 transduction of the kanamycin marker into a WT *E. coli* background and checked that, owing to their genetic proximity each corresponding cysteine-to-serine mutation were co-transduced along with the kanamycin marker. *E. coli* gene deletion used in this study originated either from the *E. coli* Keio collection of mutants (Baba et al., 2006) or were generated by λ-red linear DNA gene inactivation using pKOBEG or pKOBEGA plasmids (Chaveroche et al., 2000). Primers used to construct recombinogenic DNA fragments are listed in supplementary Table S5. P1*vir* transduction was used to transfer mutations between different strains. When required, antibiotic resistance markers flanked by two FRT sites were removed using Flp recombinase (Cherepanov and Wackernagel, 1995). Plasmids used in this study were constructed using an isothermal assembly method, Gibson assembly (New England Biolabs, Ipswich, MA, USA), using primers listed in Table S2. The integrity of all cloned fragments, mutations, and plasmids was verified by PCR with specific primers and DNA sequencing.

### Construction and characterization of *flu-lacZ* transcriptional fusions

<u>Deletion of chromosomal *lacZ* gene mutation</u> (Δ*lacIZ::Cm*) and a *endflu-lacZzeo* construct corresponding to a translational fusion between the *flu* gene encoding Ag43 autotransporter adhesin and the reporte gene *lacZ* were successively introduced by P1 *vir* phage transduction in *E. coli* WT, Δ*oxyR::Km* and *oxyR*_C25S_ strains. The On (blue) or OFF status of Agn43 in these strain were assessed by resuspending colonies in 1ml LB medium and plating dilutions on LB agar plates supplemented with 100ug.mL^−1^ of 5-bromo-4-chloro-3-indolyl-b-D-galactopyranoside (X-gal). The plates were incubated overnight at 37°C.

### Ag43 immunodetection

For each biofilm culture, the equivalent of 0,2OD_600_ units was analyzed by sodium dodecyl sulfate-10%polyacrylamide gel electrophoresis, followed by immunodetection of Ag43 using a polyclonal rabbit antiserum raised against the a-domain of Ag43 at a dilution of 1:10,000. Protein loading accuracy was verified using staining of membrane with Ponceau S red.

### Biofilm and planktonic culture conditions

Biofilms were cultivated in continuous-flow microfermentors containing a removable glass spatula (Ghigo, 2001) (see also https://research.pasteur.fr/en/tool/biofilm-microfermentors/). Sterile microfermentors were inoculated with 10^8^ bacteria from an overnight culture, and cells were allowed to attach for 1 h static at 37°C before turning on the medium flow. Microfermentors were operated with a medium flow rate of 40 mL/h (residence time 40 min) and internal bubbling of a filter-sterilized compressed gas at ~0.2 bar_g_ consisting of: (i) air for aerobic conditions, or (ii) a mix of 90% nitrogen / 5% hydrogen /5% carbon dioxide (Air Liquide) for anaerobic conditions. Biofilm cultures were grown for 48 h before protein extraction. Planktonic cultures were inoculated to an optical density of the culture at 600 nm (OD_600_) of 0.005 and grown in Erlenmeier glass flasks at 37°C, either: (i) in an aerated shaker incubator for aerobic conditions, or (ii) in a Concept 400M anaerobic workstation (Ruskinn) on a multi-position magnetic stirrer (Carl Roth) for anaerobic conditions. Planktonic cultures were grown for 24 h before protein extraction.

### Motility assay

Overnight cultures of the tested bacterial strains were spotted as a 2-μL drop of a 10^−2^ dilution onto 0.3% agar plates with 10 g/L Bacto tryptone, 5 g/L NaCl and 3 g/L agar, or with 3 g/L agar in M63B1 minimal medium. Motility plates were incubated overnight at 30°C. The distance from the edge of the inoculation spot to the edge of the growth zone was measured.

### Oxidative stress assay

Overnight cultures in LB medium were diluted to an OD_600_ of 0.05 in fresh medium and then allowed to grow to an OD_600_ of 0.1. Then, 100 μL of each culture were spread on LB or M63B1 plates. Round sterile filters were placed in the center of the plates and spotted with 25 μL of 30% H_2_O_2_. Plates were incubated at 37°C overnight. The distance from the edge of the disk to the edge of the growth zone was measured.

### Autoaggregation Assay

Isolated colonies were used to inoculate 5 mL LB medium and grown overnight (16–18 h). The OD_600_ was adjusted to 3.0 by resuspension of the cell pellet in nutrient-exhausted LB medium (supernatant obtained from respective overnight grown cultures after centrifugation), and 3 mL of each adjusted culture were transferred to 5 mL hemolysis tubes. These tubes were incubated without agitation at room temperature and the OD_600_ of the upper part of each standing tube culture was determined every hour for 8 h.

### *S*-nitrosylated and oxidized thiols biotin-switch SILAC method

#### Protein extraction

*S*-oxidized and *S*-nitrosylated peptides were analyzed by applying a cysteine biotin-switch technique, as described before (Li et al., 2015), with some modifications. First, biofilm or planktonic bacteria were lysed with trichloroacetic acid (TCA) immediately after cultivation: biofilm cells on the fermenter spatula were first resuspended in cold PBS before centrifugation to obtain the cell pellet, while 25 mL of planktonic cells were immediately centrifuged for 10 min at 8,000 x g. The cell pellet was resuspended in 0.6 mL of 20% (v/v) TCA, thus representing ~1.5-2.0 mg/mL protein content, and centrifuged at 4°C, 16,000 x g for 1 h, before washing 3 times with ice-cold acetone.

#### R-SH blocking

Following extraction, reduced thiols were blocked by resuspending the pellet containing proteins and cell debris in HENS lysis buffer, consisting of 250 mM 4-(2-hydroxyethyl)-1-piperazineethanesulfonic acid (HEPES), pH 7.7, 1 mM ethylenediaminetetraacetic acid (EDTA), 0.1 mM neocuproine, 6 M urea, 1% (w/v) N-octyl-β-D-glucopyranoside, a protease inhibitor cocktail, and supplemented with 200 mM iodoacetamide (IAM), allowing for improved saturation of reduced thiols as shown in a previous study (Shakir et al., 2017). Protein content was adjusted to 0.8 mg/mL in a volume of 400 μL and the mixture was incubated at 37°C with shaking at 2,000 rpm for 1 h in the dark in a Thermomixer (Eppendorf).

For mass spectrometry analysis, each sample was mixed with one volume of a heavy isotope-labeled reference sample after this blocking step and further processed as one sample. The reference sample consisted of a 5:5:1 mixture of planktonic extracts corresponding to non-*S*-nitrosylating (aerobic with fumarate), mildly *S*-nitrosylating (anaerobic with 300 μM nitrate) and strongly *S*-nitrosylating (anaerobic with 10 mM nitrate) conditions.

#### R-SNO ascorbate reduction and IAM-biotin labeling

Proteins were precipitated with TCA and acetone before being resuspended in HENS buffer with 10 mM ascorbate and 2 mM EZ-Link iodoacetyl-PEG2-biotin (IAM-biotin, Thermo Scientific), and incubation at 37°C with shaking at 2,000 rpm for 1 h.

At this stage, proteins were either directly analyzed by western blot or further processed for R-S-OX reduction and labeling, peptide digestion, avidin pull-down and finally identification by mass spectrometry.

#### Western blot analysis

Proteins were precipitated with TCA and acetone before being resuspended in 300 μL 50 mM ammonium bicarbonate, at 37°C with shaking at 2,000 rpm for 1 h. Then the solution was mixed with 2X Laemmli buffer without reducing agent and loaded onto SDS-PAGE, transferred to cellulose membrane by using a TranS-Blot Turbo Transfer System (Bio-Rad), labeled with avidin-HRP conjugates (eBioscience) (1:10,000 dilution in PBS with 0.05% Tween), revealed with ECL Prime chemiluminescence reagents (GE Healthcare) and visualized in a G-Box (Syngene). Total proteins were visualized from SDS-PAGE gels with Stain-free fluorescent imaging (BioRad).

#### R-SOX DTT reduction and S-S-biotin labeling

Proteins were precipitated with TCA and acetone before being resuspended in HENS buffer with 10 mM 1,4-dithiothreitol (DTT) and 0.5 mM EZ-Link HPDP-biotin (S-S-biotin, Thermo Scientific), and incubation at 60°C with shaking at 2,000 rpm for 30 min.

#### Trypsin digestion

Proteins were precipitated with TCA and acetone before being resuspended in 300 μL of 50 mM ammonium bicarbonate and digested with 1 μg/mL trypsin (Roche) at 37°C, 1,000 rpm overnight.

#### Avidin pull-down

One hundred microliters of streptavidin resin (Pierce) were loaded onto spin columns (Pierce) and washed 5 times with 700 μl PBS, before loading 300 μl trypsinized sample and incubation at 25°C, 1,000 rpm for 1 h. Unbound peptides were collected, immediately stabilized with 1% Trifluoroacetic acid and stored frozen until MS analysis. The avidin resin was washed 5 times with 600 μL bicarbonate, before eluting the R-SOX peptides with 300 μL 10 mM DTT at 37°C, 2,000 rpm for 2 h, which were collected and stabilized with 1% TFA. Finally, after washing the R-SNO peptides were eluted with 300 μL of 7 M guanidine-HCl at 95°C for 30 min.

### Mass spectrometry analysis

Mass spectrometry analysis was performed by using 5 biological replicate samples, all combined with a heavy-isotope labeled reference sample as described above. Thus, the four series of samples (aerobic biofilm, aerobic planktonic, anaerobic biofilm, anaerobic planktonic) were analyzed in independent LC-MS/MS runs. Prior to LC-MS/MS analysis, digested peptides were desalted on a C18 microcolumn (Zip-Tip, Millipore), eluted in 2 μL of 60% acetonitrile (ACN), 0.1% aqueous formic acid (FA) and added 18 μL of 0.1% FA. Each sample was concentrated (5 μL) on a C18 cartridge (Dionex Acclaim PepMap100, 5 μm, 300 μm i.d. x 5 mm) and eluted on a capillary reverse-phase column (C18 Dionex Acclaim PepMap100, 3 μm, 75 μm i.d. x 50 cm) at 220 nL/min, with a gradient of 2% to 50% of buffer B in 180 min; (buffer A: 0.1% aq. FA/ACN 98:2 (v/v); buffer B: 0.1% aq. FA/ACN 90:10 (v/v)), coupled to a quadrupole Orbitrap mass spectrometer (Q Exactive, ThermoFisher Scientific) using a top 10 data-dependent acquisition MS experiment: 1 survey MS scan (400-2,000 m/z; resolution 70,000) followed by 10 MS/MS scans on the 10 most intense precursors (dynamic exclusion of 30 s, resolution 17,500).

#### Data analysis

Database search using MaxQuant (version 1.3.0.5) on SwissProt database (2017_05) was performed with methionine oxidation, cysteine carbamidomethylation (IAM) and biotinylation (iodoacetyl-PEG2-biotin, IAM-biotin) as variable modifications, with the following conditions: first search error tolerance 20 ppm, main error tolerance 6 ppm, MS/MS error tolerance 20 ppm, FDR 1%. A fold change was calculated for each condition: internal standard/aerobic biofilm, internal standard/aerobic planktonic, internal standard/anaerobic biofilm and internal standard/anaerobic planktonic. Quantitative profiles were generated by reprocessing the data from MaxQuant evidence file. Protein expression profiles were estimated using cysteine free peptides only. We considered peptides carrying only one cysteine. Cysteine containing peptides were classified according to their oxidation state (IAM labeled for reduced, IAM-biotin for S-NO, unmodified for S-OX). When a given cysteine residue was detected in different peptide forms (miss cleavages or oxidized methionine containing peptides), the features were aggregated using the median value of the ratios. The fold changes of cysteine-containing peptide were normalized to the related protein fold changes in order to avoid quantification bias induced by the differential protein expression. The null hypothesis test was performed by comparing the modified peptide fold change to the associated protein fold change. The normalized ratios (cysteine fold changes and protein fold changes) with the corresponding calculated variances were used to perform Student’s test between biofilm and planktonic conditions. P-values < 0.05 and fold changes > 1.75 were chosen as significant thresholds. Proteins detected in 4 or 5 replicates and absent in the other conditions were labeled as “biofilm only” or “planktonic only”. The same procedure was used for the cysteine-containing peptides, considering only species associated to proteins detected in both conditions.

Proteins with significant fold changes were used to create a dataset of “upregulated” and “downregulated” entries. The datasets were uploaded in STRING (Szklarczyk et al., 2019) to perform protein network analyses and gene ontology enrichments, using K-means clustering, which produced 6 clusters.

## ACKNOWLEDGEMENTS

We thank Rebecca Stevick for critical reading of the manuscript. This work was supported by the Institut Pasteur, the French Government’s *Investissement d’Avenir* program: Laboratoire d’Excellence ‘Integrative Biology of Emerging Infectious Diseases (LabEx IBEID) (grant n°. ANR-10-LABX-62-IBEID to J.M.G.), the *Fondation pour la Recherche Médicale* (grant n°. DEQ20180339185 to J.M.G.). Mass spectrometry equipment was subsidized by Conseil Régional d’Île-de-France (Sesame 2010 N°10022268). N.B. was the recipient of a LabEx IBEID post-doctoral fellowship.

## DATA AVAILABLITY STATEMENT

Data are available via ProteomeXchange with identifier PXD020249.

## AUTHOR CONTRIBUTIONS

N.B., G.C., S.L., J.V. and J.-M.G. designed the experiments. N.B., G.C. and S.L. performed the experiments. N.B., G.C. and J.-M.G. analyzed the data and wrote the paper with significant contribution from S.L. and J.V. All authors have read and approved the manuscript.

## SUPPLEMENTARY INFORMATION

### SUPPLEMENTARY FIGURES (supplementary Figures S1–S8)

**Supplementary Figure S1.**
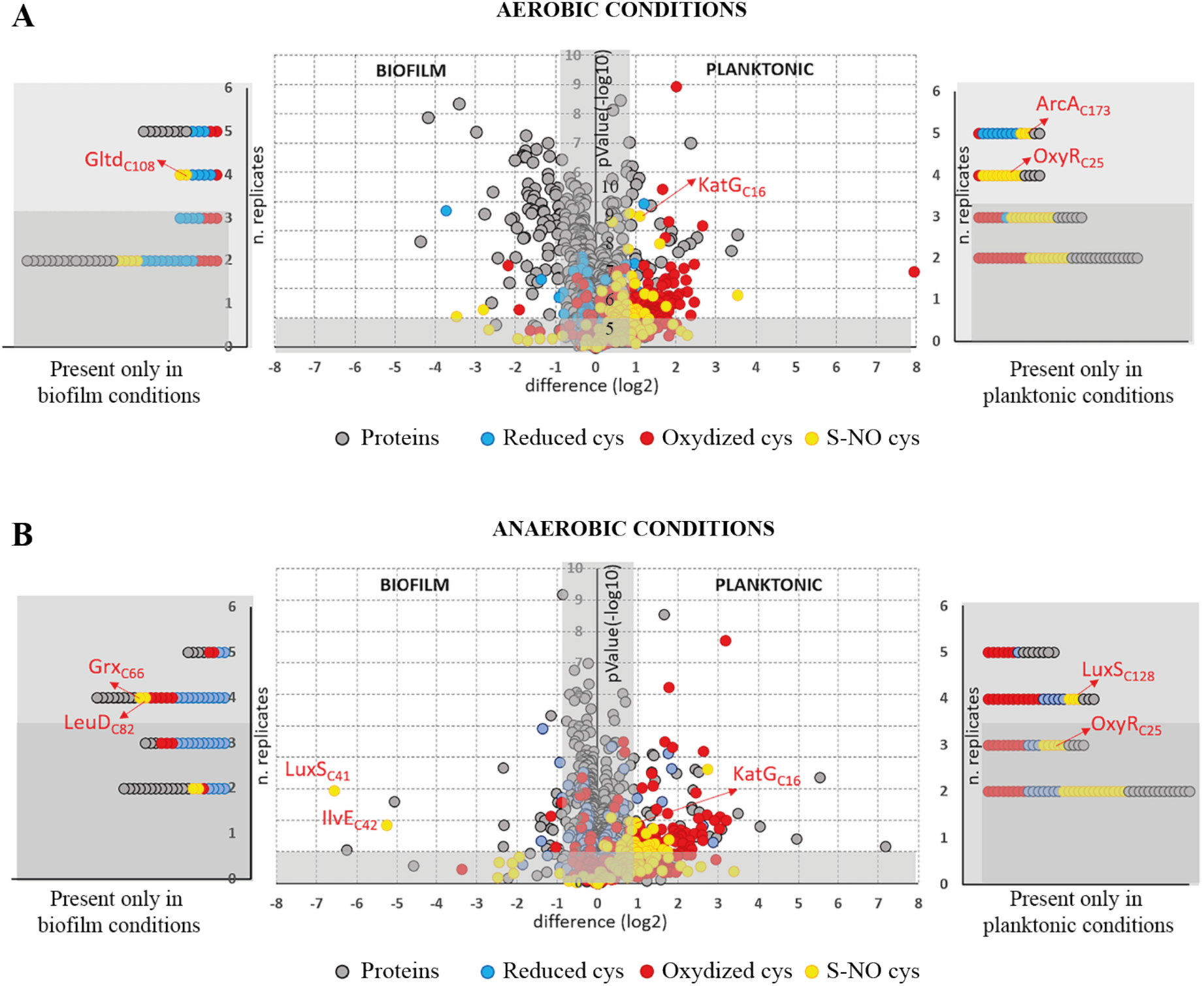
Volcano plots combining the global dataset of the cysteine redox proteomics analysis of aerobic (panel A) and anaerobic (panel B) *E. coli* biofilm and planktonic cultures. Proteins appear in grey, reduced cysteines in blue, reversibly oxidized S-OX cysteines in red, *S*-nitrosylated S-NO cysteines in yellow. The lateral panels show the features detected specifically to one lifestyle. The dark grey areas delimit the non-significant data (thresholds as described in the method section).

**Supplementary Figure S2.**
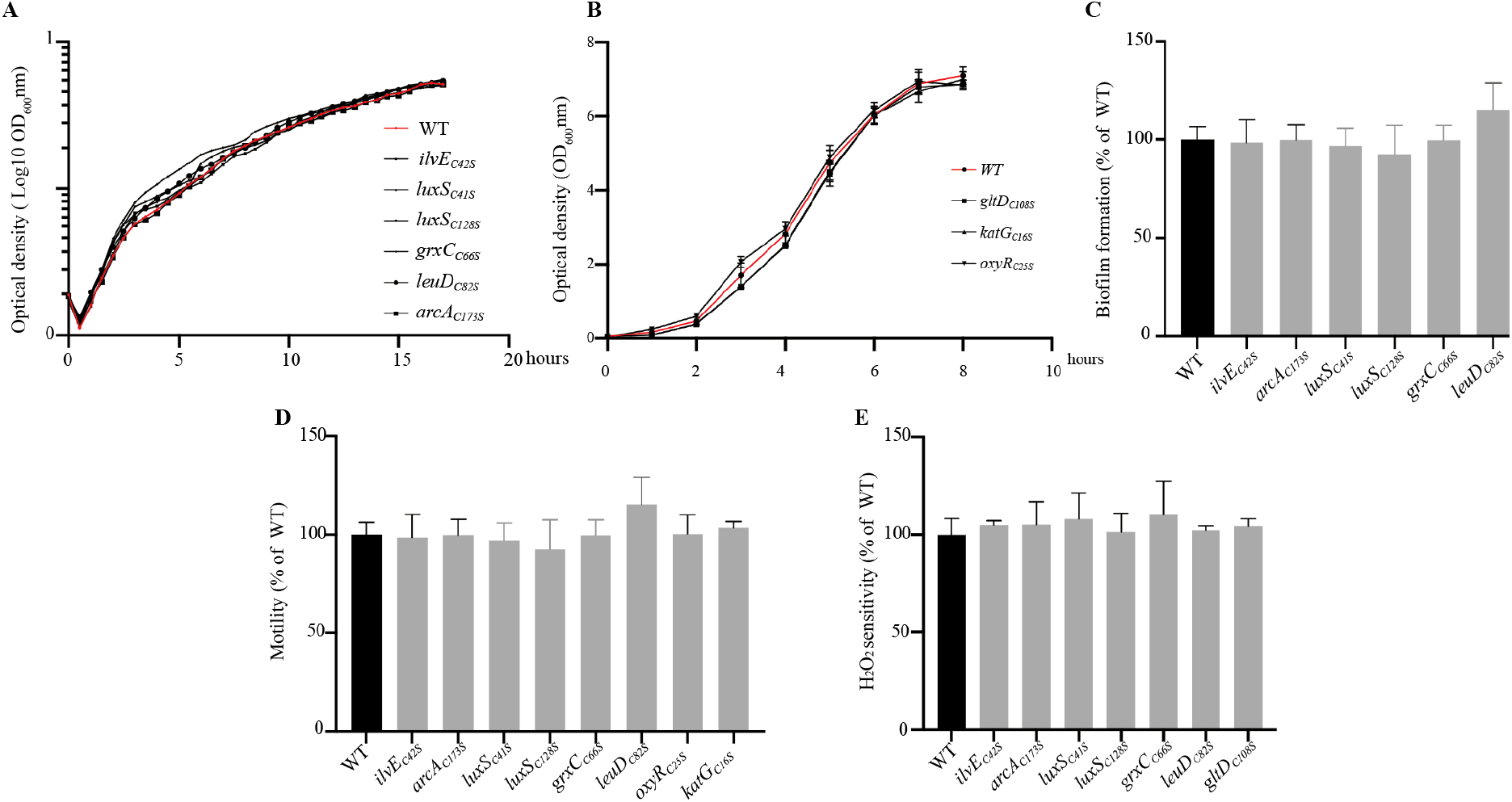
Phenotypic analysis of mutants in genes corresponding to proteins with differentially *S-nitrosylated* cysteines. **A**: Growth kinetics were assessed in aerobic planktonic conditions in LB medium for *E. coli* WT (MG1655 F’*tet*), *grxC*_C66S_, *leuD*_C82S_, *ilvE*_C42S_, *luxS*_C41S_, *luxS*_C128S_, *arcA*_C173S_ mutants. **B**: Growth kinetics were assessed in aerobic planktonic conditions in LB medium for *oxyR*_C25S_, *katG*_C16S_ and *gltD*_C108S_ mutants and WT strain. **C**: Biofilm biomass formation in microfermentors after 24 h in LB medium for *grxC*_C66S_, *leuD*_C82S_, *ilvE*_C42S_, *luxS*_C41S_, *luxS*_C128S_, *luxS*_C128S_ mutants and WT strains. The level of biofilm formed by the WT strain was set to 100%. **D**: Motility assays on 0.3% agar plates, incubated overnight at 30°C for *grxC*_C66S_, *leuD*_C82S_, *ilvE*_C42S_, *luxS*_C41S_, *luxS*_C128S_, *arcA*_C173S_, *oxyR*_C25S_, *katG*_C16S_ mutants and WT strains. The diameter of the growth zone was measured and set to 100% for the WT strain. **E**: Sensitivity to H_2_O_2_ oxidative stress for *grxC*_C66S_, *leuD*_C82S_, *ilvE*_C42S_, *luxS*_C41S_, *luxS*_C128S_, *arcA*_C173S_ and *gltD*_C108S_ mutants and WT strains. The distance from the edge of the disk to the edge of the growth zone was measured and was set to 100% for the WT strain. All experiments were performed in triplicate, mean values are reported and error bars represent standard deviations.

**Supplementary Figure S3.**
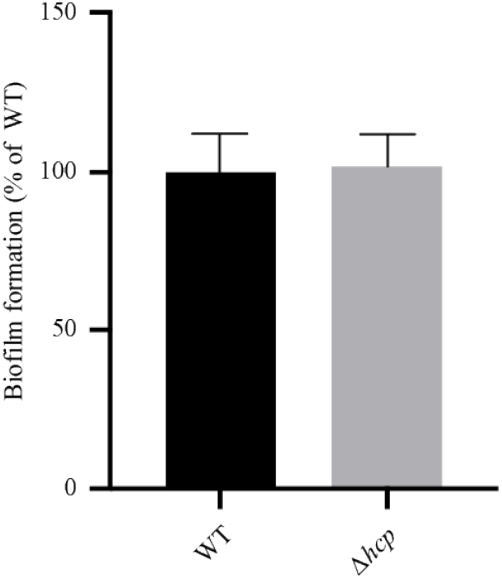
Δ*hcp* mutant biofilm formation. Biofilms were cultivated in microfermentors in LB medium for 24 h. The level of biofilm formed by the *E. coli* WT (MG1655 F’*tet*) strain was set to 100%. Data are the means from three independent experiments.

**Supplementary Figure S4.**
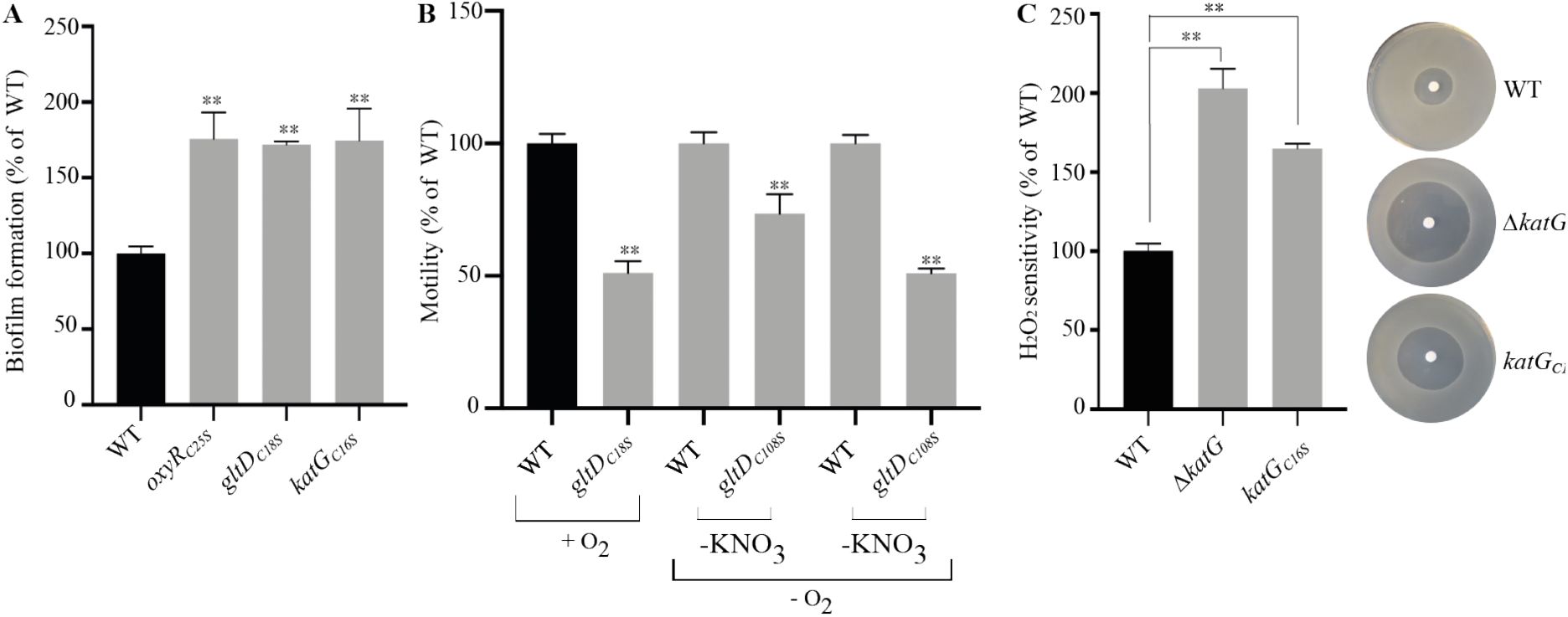
Phenotypes associated with OxyR, KatG and GltD *S*-nitrosylation in M63B1 minimal medium. A: Biofilms of *E. coli* WT, *oxyR*_C25S_, *katG*_C16S_ and *gltD*_C108S_ mutants, were grown in continuous-flow microfermentors for 48 h in M63B1 glucose medium before quantifying the biofilm biomass. The level of biofilm formed by the WT strain was set to 100%. Data are the means from three experiments. **B**: The motility of *E. coli* WT (MG1655 F’*tet*) and *gltD*_C108S_ mutant were compared in M63B1 glucose medium in aerobic conditions and in anaerobic conditions in presence of KNO_3_. Assays were performed on 0.3% agar plates and incubated overnight at 30°C. The diameter of the growth zone was measured and set to 100% for each WT strain. **C**: Sensitivity to H_2_O_2_ oxidative stress of a *katG*_C16S_ mutant compared to the WT. The distance of growth inhibition from the edge of the disk to the edge of the growth zone was measured and was set to 100% for the WT strain. All experiments were performed in triplicate, mean values are reported and error bars represent standard deviations. ** p ≤ 0.05, *** p ≤ 0.01.

**Supplementary Figure S5.**
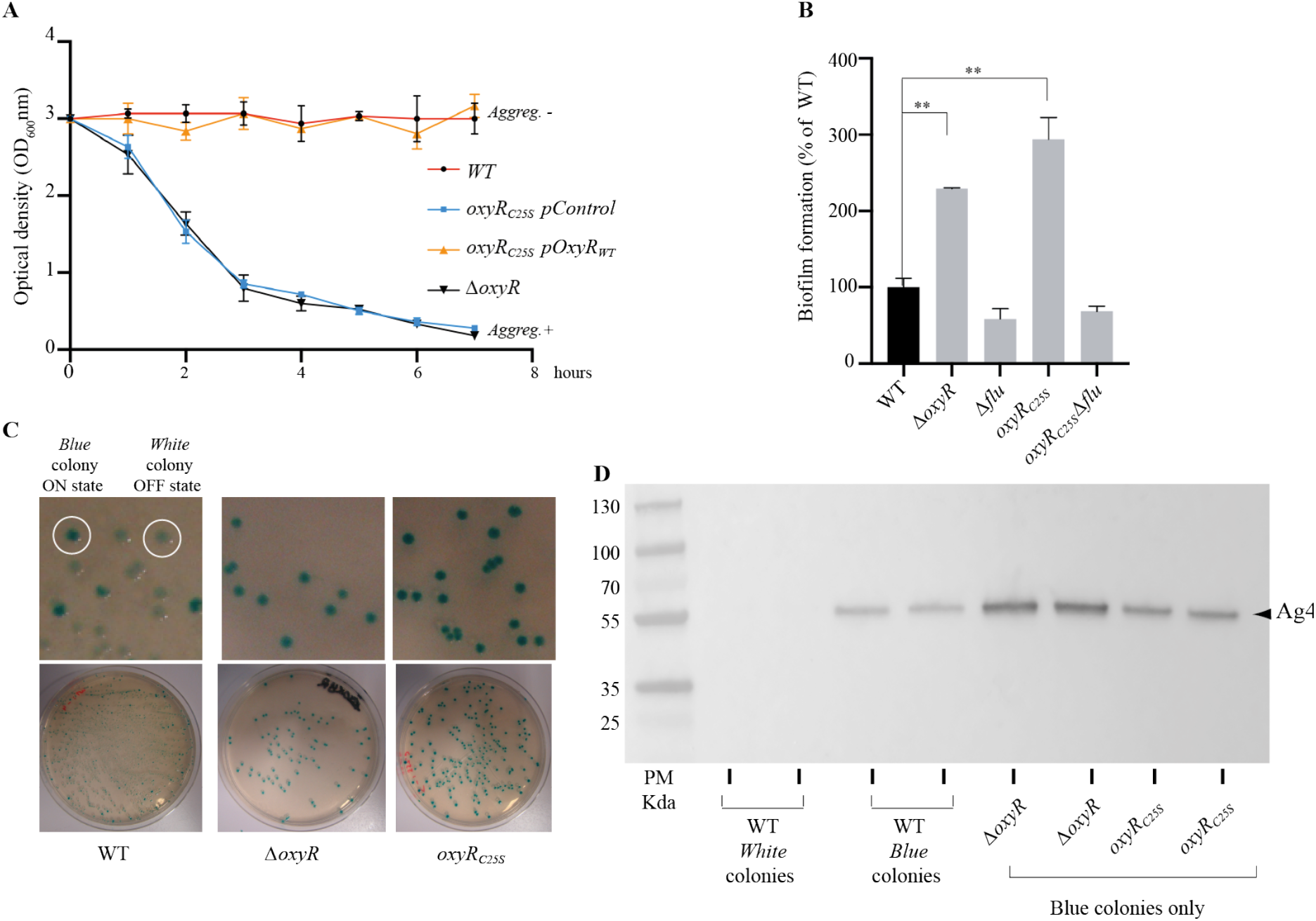
Impairing *S*-nitrosylation status of OxyRC25 cysteine affects aggregation and *flu*-dependent biofilm formation. **A**: Cell aggregation kinetics for *oxyR*_C25S_ mutant, complemented with corresponding plasmid-based allele or empty vector. **B**: Biofilm formation of *oxyR*_C25S_ and Δ*flu* mutants. The level of biofilm formed by the WT strain was set to 100%. All experiments were performed in triplicate, mean values are reported and error bars represent standard deviations. ** p ≤ 0.05, *** p ≤ 0.01. **C**: *flu*(=*ag43*)-*lacZ* transcriptional fusions in a WT, Δ*oxyR* and *oxyR*_C25S_ background plated on LB agar + X-gal plates: Blue colony: ag43 expresion in an ON state. White colony: ag43 expresion in an OFF state **D:**Immunodetection of Ag43 in biofilm cultures of 2 white or blue colonies of WT, 2 blue colonies of Δ*oxyR* and two blue colonies of *oxyR*_C25S_ using an anti-Ag43 polyclonal antibody.

**Supplementary Figure S6.**
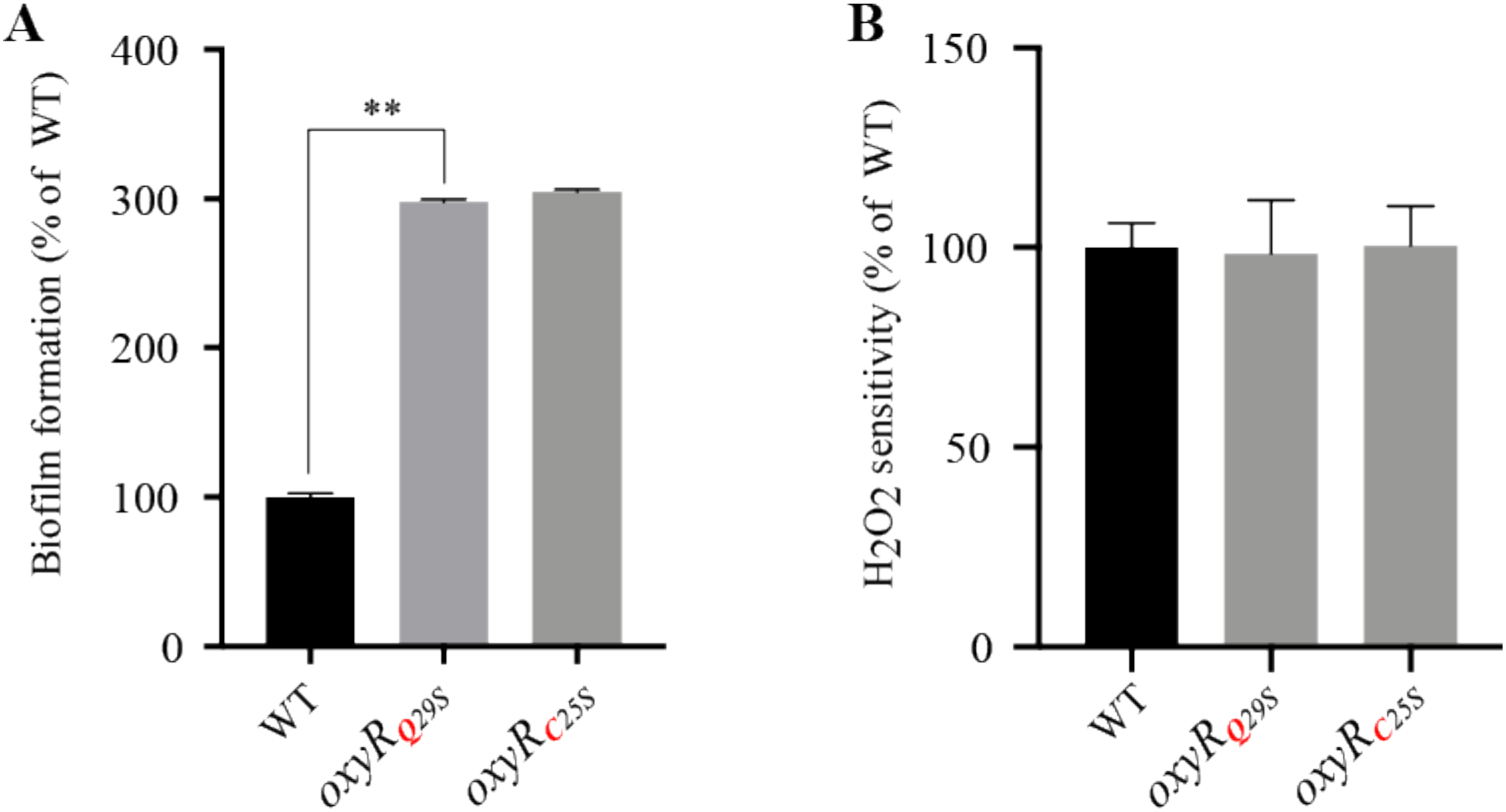
Impairing OxyR DNA binding site by mutating increases biofilm formation but has no impact on *E. coli* H_2_O_2_ sensitivity. A: Biofilms of *E. coli* WT, *oxyR*_C25S_ and *oxyR*_Q29S_ mutants were grown in continuous flow microfermentors for 24 h in LB medium before quantifying the biofilm biomass. The level of biofilm formed by the WT strain was set to 100%. B: Sensitivity to H_2_O_2_ oxidative stress of *oxyR*_C25S_ and *oxyR*_Q29S_ mutants compared to the WT. The distance of growth inhibition from the edge of the disk to the edge of the growth zone was measured and was set to 100% for the WT strain. All experiments were performed in triplicate, mean values are reported and error bars represent standard deviations. ** P ≤ 0.05, *** P ≤ 0.01.

**Supplementary Figure S7:**
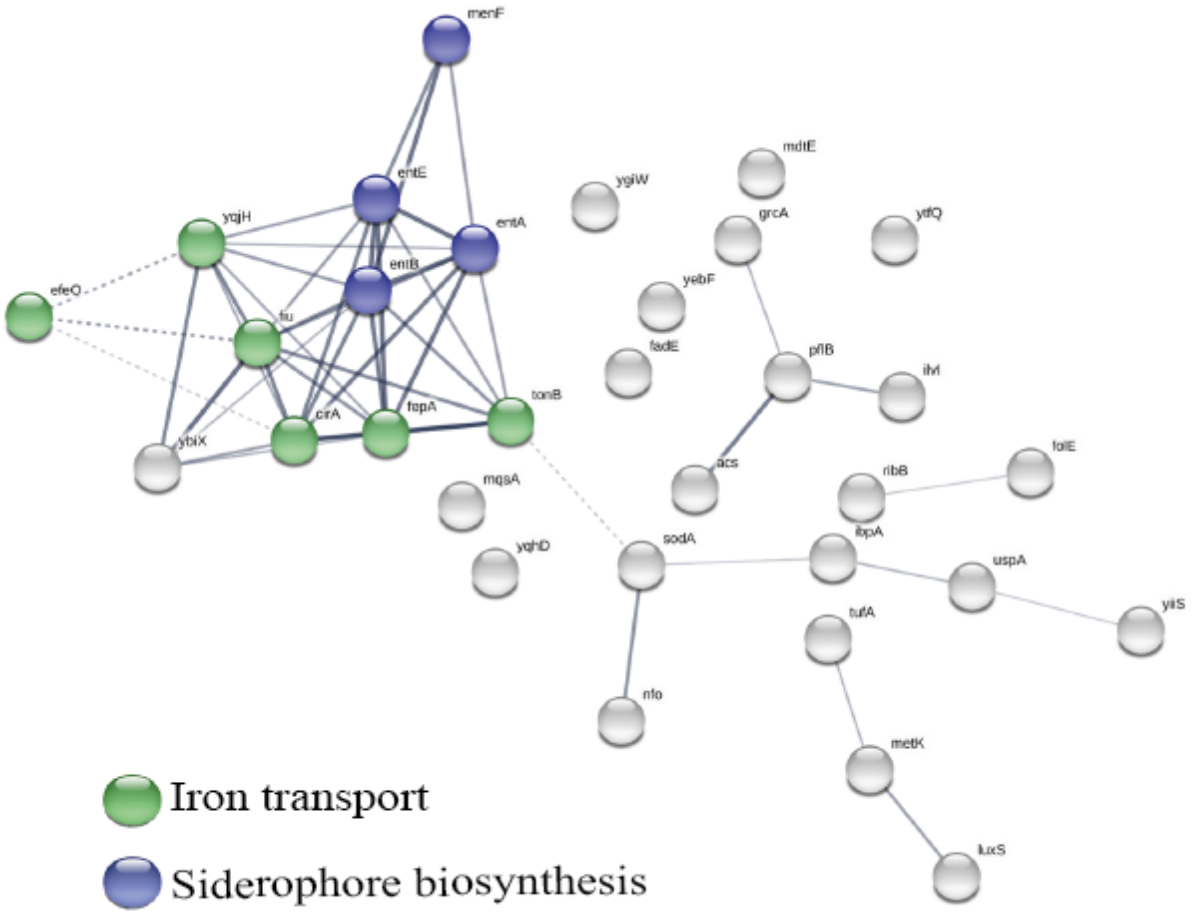
STRING network analysis of proteins downregulated in biofilms. One significative cluster was obtained by k-means clustering. The proteins belonging to the cluster are involved in siderophore biosynthesis (blue spheres) and iron transport (green spheres).

**Supplementary Figure S8:**
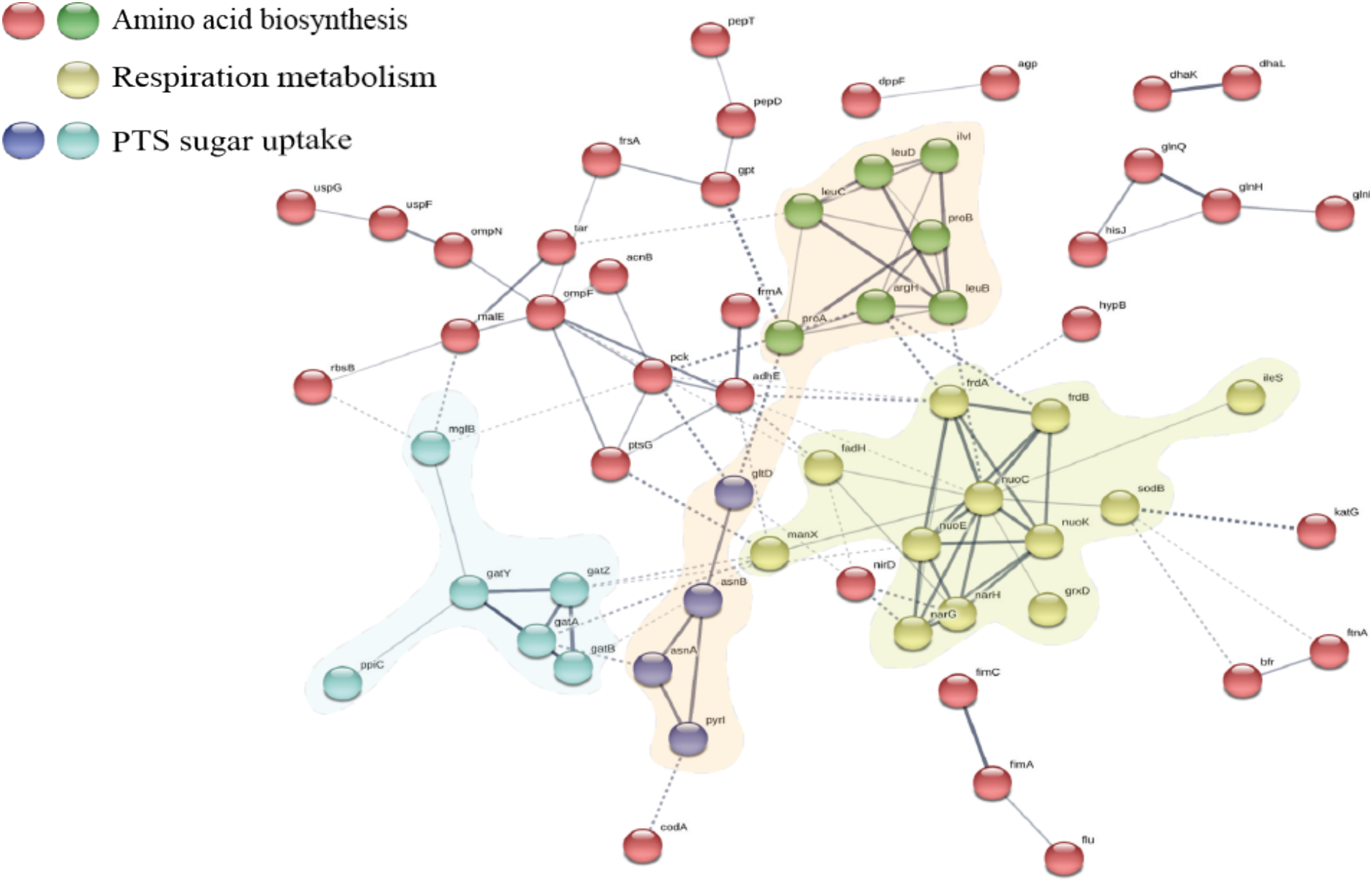
STRING network analysis of proteins upregulated in biofilms. Three significative clusters were obtained by k-means clustering. Proteins in the cluster represented in yellow are involved in the respiratory chain. It is characterized by 11 oxidoreductase proteins, 7 of which (NuoC, NuoE, NuoK, NarG, NarH, FrdA, FrdB) are involved in respiration. Protein clusters represented in green and dark blue are involved in amino acid biosynthesis (LeuA, LeuC, LeuD, ProA, ProB, IlvI, ArgH) and (AsnA, AsnB, GltD, PyrL). Proteins in the light blue cluster belong to the PEP group translocation mechanism for sugar uptake and consists of 6 proteins, 4 of which (GatA, GatB, GatY, GatZ) belong to a PTS system involved in galactitol uptake.

### SUPPLEMENTARY TABLES (supplementary Tables S1–S5)

**Supplementary Table S1.** List of peptides differentially reduced, *S*-oxidized and *S*-nitrosylated in biofilm *vs* planktonic *E. coli* cultures under aerobic conditions, by using the biotin-switch SILAC method. PO, planktonic with O_2_; BO, biofilm with O_2_. Content description in the first sheet of the Excel spreadsheet.

**Supplementary Table S2.** List of peptides differentially reduced, *S*-oxidized and *S*-nitrosylated in biofilm *vs* planktonic *E. coli* cultures under anaerobic conditions, by using the biotin-switch SILAC method. PN, planktonic without O_2_; BN, biofilm without O_2_. Content description in the first sheet of the Excel spreadsheet.

**Supplementary Table S3.** List of proteins identified to be differentially expressed in biofilm *vs* planktonic *E. coli* cultures by using the biotin-switch SILAC method. PO, planktonic with O_2_; PN, planktonic without O_2_; BO, biofilm with O_2_; BN, biofilm without O_2_. Content description in the first sheet of the Excel spreadsheet.

**Supplementary table S4.**
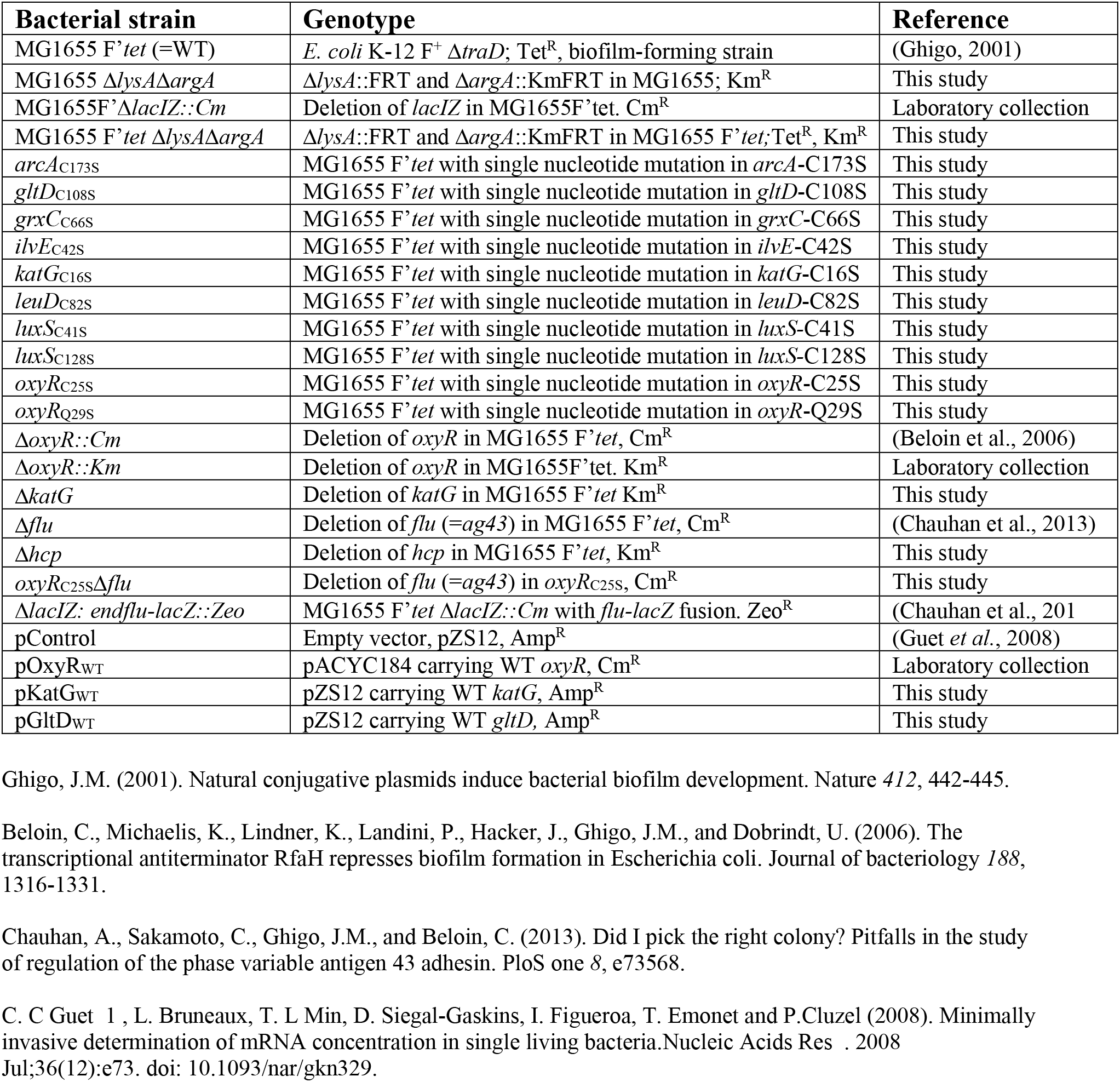
Bacterial strains and plasmids used in this study

| Bacterial strain | Genotype | Reference |
| --- | --- | --- |
| MG1655 F'tet (=WT) | *E. coli* K-12 F<sup>+</sup> Δ*traD*; Tet<sup>R</sup>, biofilm-forming strain | (Ghigo, 2001) |
| MG1655 Δ*lysA*Δ*argA* | Δ*lysA*::FRT and Δ*argA*::KmFRT in MG1655; Km<sup>R</sup> | This study |
| MG1655F'Δ*lacIZ*::*Cm* | Deletion of *lacIZ* in MG1655F'tet. Cm<sup>R</sup> | Laboratory collection |
| MG1655 F'tet Δ*lysA*Δ*argA* | Δ*lysA*::FRT and Δ*argA*::KmFRT in MG1655 F'tet;Tet<sup>R</sup>, Km<sup>R</sup> | This study |
| *arcA*<sub>C173S</sub> | MG1655 F'tet with single nucleotide mutation in *arcA*-C173S | This study |
| *gltD*<sub>C108S</sub> | MG1655 F'tet with single nucleotide mutation in *gltD*-C108S | This study |
| *grxC*<sub>C66S</sub> | MG1655 F'tet with single nucleotide mutation in *grxC*-C66S | This study |
| *ilvE*<sub>C42S</sub> | MG1655 F'tet with single nucleotide mutation in *ilvE*-C42S | This study |
| *katG*<sub>C16S</sub> | MG1655 F'tet with single nucleotide mutation in *katG*-C16S | This study |
| *leuD*<sub>C82S</sub> | MG1655 F'tet with single nucleotide mutation in *leuD*-C82S | This study |
| *luxS*<sub>C41S</sub> | MG1655 F'tet with single nucleotide mutation in *luxS*-C41S | This study |
| *luxS*<sub>C128S</sub> | MG1655 F'tet with single nucleotide mutation in *luxS*-C128S | This study |
| *oxyR*<sub>C25S</sub> | MG1655 F'tet with single nucleotide mutation in *oxyR*-C25S | This study |
| *oxyR*<sub>Q29S</sub> | MG1655 F'tet with single nucleotide mutation in *oxyR*-Q29S | This study |
| Δ*oxyR*::*Cm* | Deletion of *oxyR* in MG1655 F'tet, Cm<sup>R</sup> | (Beloin et al., 2006) |
| Δ*oxyR*::*Km* | Deletion of *oxyR* in MG1655F'tet. Km<sup>R</sup> | Laboratory collection |
| Δ*katG* | Deletion of *katG* in MG1655 F'tet Km<sup>R</sup> | This study |
| Δ*flu* | Deletion of *flu* (=ag43) in MG1655 F'tet, Cm<sup>R</sup> | (Chauhan et al., 2013) |
| Δ*hcp* | Deletion of *hcp* in MG1655 F'tet, Km<sup>R</sup> | This study |
| *oxyR*<sub>C25S</sub>Δ*flu* | Deletion of *flu* (=ag43) in *oxyR*<sub>C25S</sub>, Cm<sup>R</sup> | This study |
| Δ*lacIZ*::*endflu-lacZ*::*Zeo* | MG1655 F'tet Δ*lacIZ*::*Cm* with *flu-lacZ* fusion. Zeo<sup>R</sup> | (Chauhan et al., 2013) |
| pControl | Empty vector, pZS12, Amp<sup>R</sup> | (Guet et al., 2008) |
| pOxyR<sub>WT</sub> | pACYC184 carrying WT *oxyR*, Cm<sup>R</sup> | Laboratory collection |
| pKatG<sub>WT</sub> | pZS12 carrying WT *katG*, Amp<sup>R</sup> | This study |
| pGltD<sub>WT</sub> | pZS12 carrying WT *gltD*, Amp<sup>R</sup> | This study |

Ghigo, J.M. (2001). Natural conjugative plasmids induce bacterial biofilm development. *Nature* 412, 442-445.
Beloin, C., Michaelis, K., Lindner, K., Landini, P., Hacker, J., Ghigo, J.M., and Dobrindt, U. (2006). The transcriptional antiterminator RfaH represses biofilm formation in *Escherichia coli*. *Journal of bacteriology* 188, 1316-1331.
Chauhan, A., Sakamoto, C., Ghigo, J.M., and Beloin, C. (2013). Did I pick the right colony? Pitfalls in the study of regulation of the phase variable antigen 43 adhesin. *PloS one* 8, e73568.
C. C Guet 1, L. Bruneaux, T. L Min, D. Siegal-Gaskins, I. Figueroa, T. Emonet and P.Cluzel (2008). Minimally invasive determination of mRNA concentration in single living bacteria. *Nucleic Acids Res* . 2008 Jul;36(12):e73. doi: 10.1093/nar/gkn329.

**Supplementary Table S5:**
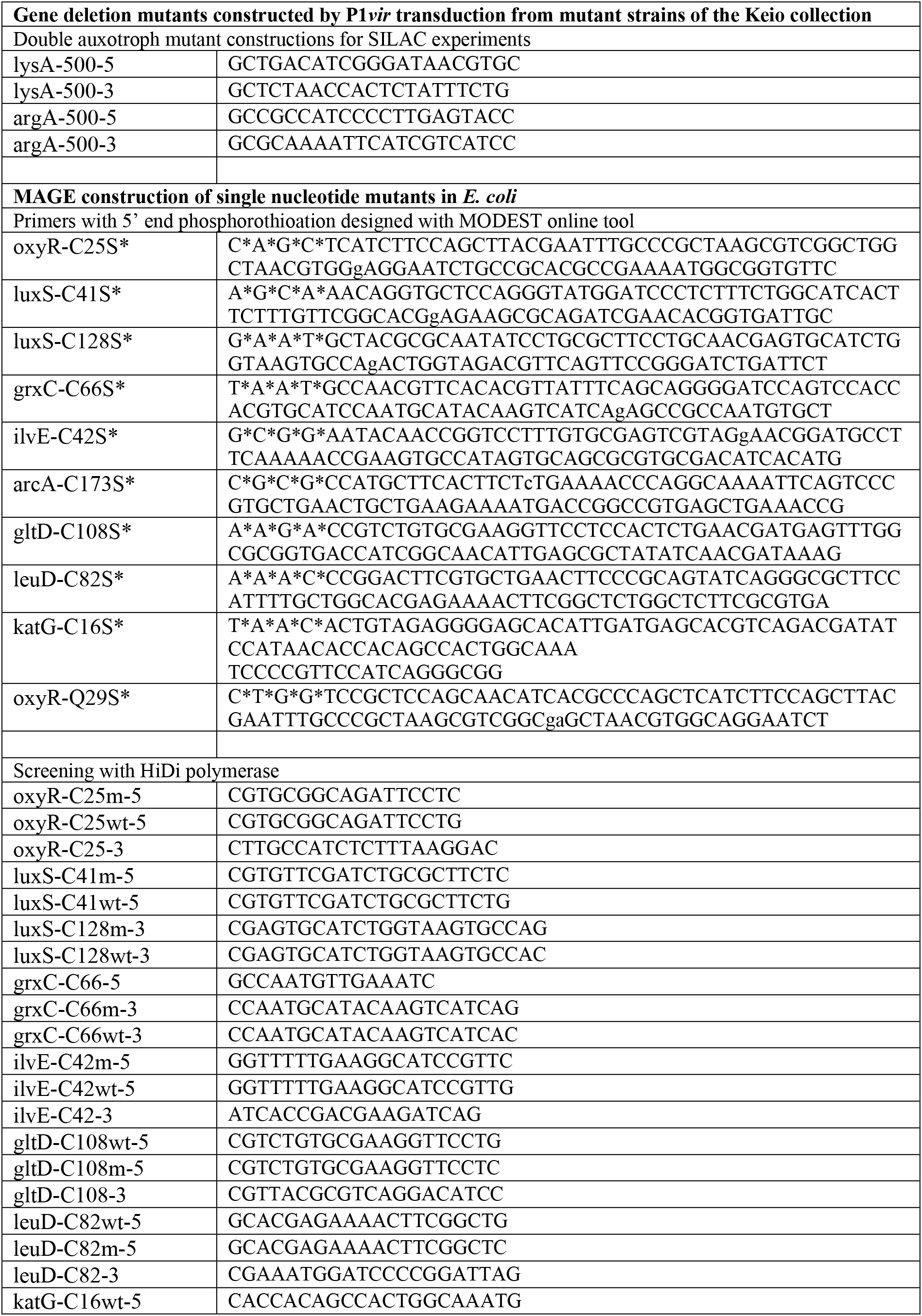
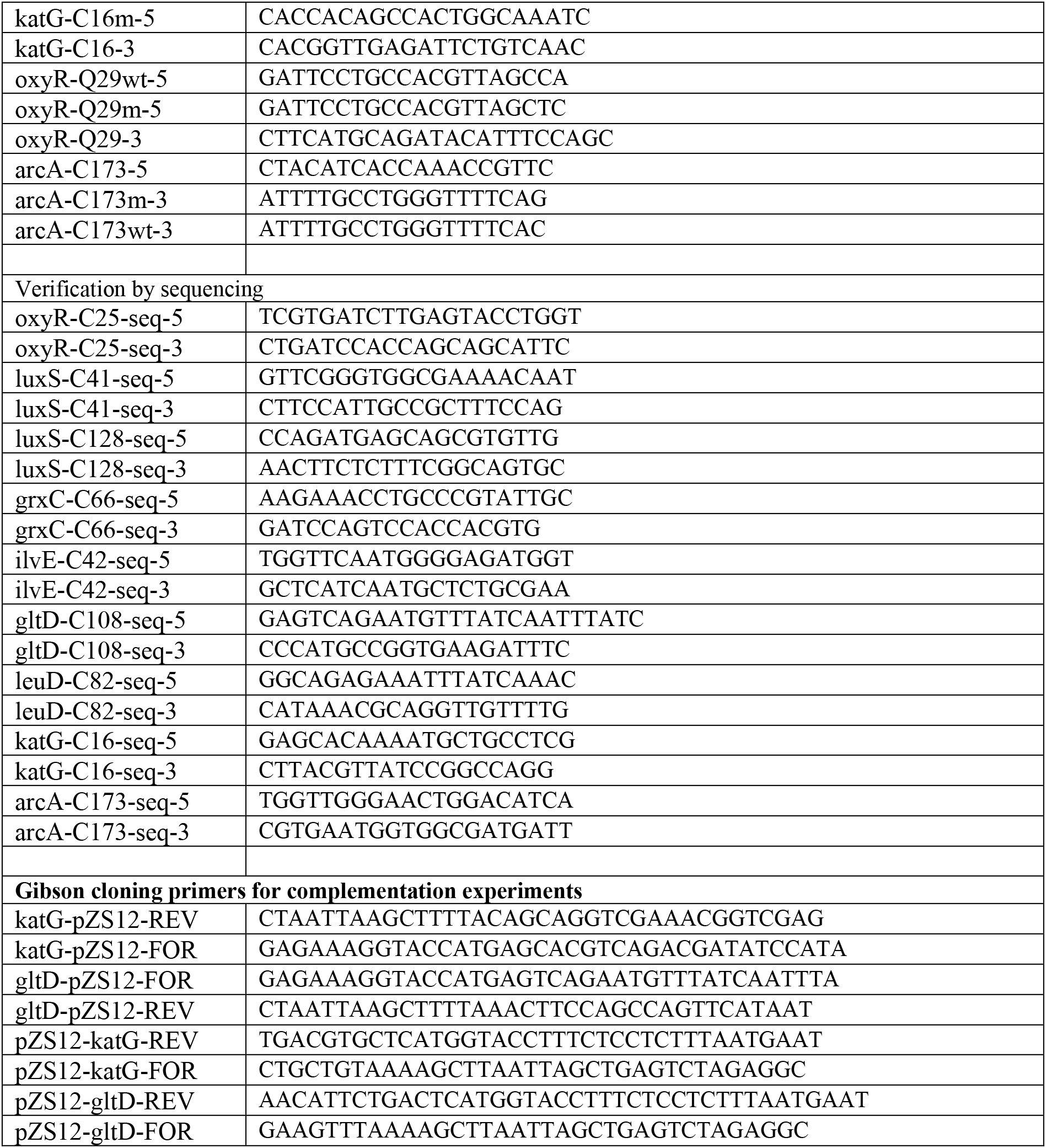
List of primers used in this study

